# Statistical inference of the cellular origin of chronic myeloid leukemia using a discrete-parameter ABC–PMC framework

**DOI:** 10.1101/2025.10.17.683025

**Authors:** Sylvie Vande Velde, Céline Engelbeen, Ilaria S Pagani, David M Ross, Sophie Hautphenne

## Abstract

Chronic myeloid leukemia (CML) arises from the *BCR::ABL1* fusion gene, but the exact stage of cellular differentiation at which the first leukemic cell emerges remains uncertain. We develop a stochastic 27-compartment model of hematopoiesis (blood cell development) using a continuous-time multitype branching process to capture the dynamics of both healthy and cancer cells. To infer the origin of CML, we develop a discrete-parameter Approximate Bayesian Computation – Population Monte Carlo (ABC–PMC) algorithm, tailored to estimate the posterior distribution for the stage of differentiation at which the first cancer cell appeared. Applied to patient data, our method consistently identifies the stem cell compartment as the most likely source of CML. These findings improve understanding of disease initiation and demonstrate the power of discrete-parameter ABC–PMC for statistical inference in complex biological systems.

**Author summary:** Chronic myeloid leukemia is a blood cancer that begins when a genetic change called the *BCR::ABL1* fusion gene appears in one cell. Although this disease has been widely studied, some questions remain, particularly about the exact stage of blood cell development at which the first cancer cell arises. In our study, we build a stochastic model based on a biological hematopoiesis model that represents how blood cells grow and mature through many stages, from stem cells to fully developed white blood cells. Using this mathematical model, we develop a statistical approach that can infer from patient data where in this hierarchy the disease most likely began. When we apply the method to clinical data from patients with chronic myeloid leukemia, it consistently points to the stem cell stage as the most probable origin. By linking biological data with mathematical modelling, our work offers new insight into how this cancer starts and shows how quantitative approaches can help answer questions that are difficult to test experimentally.

## 1 Introduction

Chronic myeloid leukemia (CML) accounts for approximately 15% of newly diagnosed leukemias in adults [1]. The reported annual incidence rate of CML ranges from 0.4 to 1.75 cases per 100,000 people [2]. CML is very rare in children, and the median age at diagnosis is around 64 years. The disease affects men more frequently than women, with approximately 60% of cases occurring in men and 40% in women [3].

CML is defined by the presence of the *BCR::ABL1* fusion gene. In over 95% of patients, CML is associated with a chromosomal translocation between chromosome 9 and chromosome 22, resulting in two abnormal chromosomes. This translocation fuses the *BCR* gene from chromosome 22 with the *ABL1* gene from chromosome 9. *BCR::ABL1* forms on the derivative chromosome 22, known as the *Philadelphia chromosome*, named after the city where it was first identified in 1960 by Peter Nowell and David Hungerford [4].

Recent studies in mouse models [5], human cells in vitro [6], and analyses of age-specific CML incidence [7], suggest that *BCR::ABL1* expression alone may be sufficient to initiate the disease. The *BCR::ABL1* fusion gene codes for an oncogenic tyrosine kinase, which plays a role in a cascade of proteins that control the cell cycle [8]. This kinase remains constitutively active, even in the absence of external signals, leading to impaired differentiation of immature blood cells and uncontrolled proliferation of white blood cells of myeloid lineage. This dysregulation causes characteristic features of CML, such as leukocytosis and enlargement of the spleen.

In addition, the *BCR::ABL1* kinase modulates the response to DNA damage, making the cells more susceptible to developing additional genetic abnormalities [9]. Because of this genetic instability, CML can progress without treatment from the chronic phase, which is the stage at which most patients are diagnosed, to a blastic phase that is almost universally fatal without transplantation. In this paper, we focus exclusively on the chronic phase.

Effective treatment with a tyrosine kinase inhibitor (TKI) prevents the disease from progressing. The first TKI, imatinib, was developed in the late 1990s, replacing earlier, less effective therapies [10]. This made CML the first cancer that could be effectively treated with oral, targeted therapy. However, some patients either do not respond to imatinib or develop resistance during treatment. As a result, second-generation TKIs such as nilotinib, dasatinib, and bosutinib were developed [11]. These therapies have increased the 10-year overall survival of CML patients to about 90%, which is comparable to that of the age-matched general population [12].

Various mathematical models, which aim to reproduce the spread of cancer cells in the blood of patients over time, have contributed to a better understanding of CML; see for example [13–16]. Most of these models are deterministic and consider different stages of cell differentiation, reflecting the hierarchical organisation of the hematopoietic (blood) system. At the top of this hierarchy are the *hematopoietic stem cells* (HSCs), located predominantly in the bone marrow. These cells differentiate to mature blood cells through successive stages involving progenitor and intermediate cell types. Mathematical models of CML typically consider between four [13] and 32 [14] differentiation stages. They address a range of biological issues, including the impact of *BCR::ABL1* expression on cell dynamics, the effect of treatment on cancer cells, the study of treatment discontinuation and the identification of factors reducing the risk of relapse, the investigation of drug resistance, and disease progression to blast crisis; see [15] for a review of these models.

To our knowledge, existing models have not yet explicitly addressed the question of the cellular differentiation stage at which the first cancer cell is most likely to arise. Most models assume that the disease originates from a hematopoietic stem cell known as a leukemia stem cell (LSC) [15]. However, recent evidence suggests that LSCs may also arise from progenitor cells rather than exclusively from HSCs [17].

To explore this question further, we develop a 27-compartment model of the hematopoietic system, where each compartment represents a distinct stage of cell differentiation. Our model adopts a stochastic approach, using a continuous-time multitype branching process called *Markovian binary tree* (MBT) to describe the dynamics of both healthy and cancer cells under different parameter sets. This stochastic framework captures the inherent randomness of cell populations, which is particularly important in early compartments where cell numbers are small. The model can generate simulations of the spread of healthy or cancerous cells across compartments over time — that is, the evolution of the disease in “virtual patients” — as well as analytical expressions for key quantities such as the expected number of cells in each compartment over time. It is tailored to incorporate current biological knowledge and can replicate clinical observations, including the elevated daily production of white blood cells in CML patients.

To infer the stage of cellular differentiation at which the first cancer cell appeared, we integrate our stochastic CML model into a discrete-parameter version of the Approximate Bayesian Computation–Population Monte Carlo (ABC–PMC) algorithm [18, 19]. ABC–PMC has previously been applied to infer parameters of an MBT in [20], in the context of phylogenetic trees, where the parameters of interest were continuous phase-specific speciation (birth) and extinction (death) rates. Here, we develop a discretised version of ABC–PMC to account for the fact that differentiation stages are naturally indexed by integers. We demonstrate that this adaptation of ABC–PMC can estimate the posterior distribution of discrete parameters from observational data and we apply it to the specific question of CML initiation. This allows us to compute both the posterior distribution and the maximum a posteriori estimate for the likely origin of the first cancer cell, based on cancer cell counts at diagnosis. We apply our approach to data from a cohort of chronic-phase CML patients in the Australian Leukemia and Lymphoma Group CML_9_ TIDEL-II clinical trial treated with frontline imatinib [21]. Across all patients, our results consistently point to the stem cell compartment as the most likely origin of CML, providing new quantitative support for the hypothesis that the disease may begin in the earliest hematopoietic stages.

The remainder of the paper is structured as follows. Section 2 describes the process of hematopoiesis for both healthy and cancer cells, along with the associated parameter values. We introduce our stochastic model in Section 3. In Section 4, we present our discrete version of the ABC–PMC algorithm, following a detailed explanation of the continuous-parameter case. Section 5 reports numerical results, first using simulated data and then applying the method to real patient data.

## 2 Models

### 2.1 Hematopoiesis model and biological parameters

We start by presenting a biological model of hematopoiesis, which describes the progression of cells from hematopoietic stem cells to fully differentiated blood cells. Depending on the choice of parameter values, this model can represent either healthy or cancer cell dynamics. The specific parameter values used in each case are detailed in Section 2.1.1 (healthy cells) and Section 2.1.2 (cancer cells).

#### Hematopoiesis model

The hematopoiesis model described by Dingli *et al*. [14] represents the progression of cells from stem cells in the bone marrow to mature blood cells using a compartmental structure. Compartment 1 corresponds to hematopoietic stem cells, and the final compartment (27 in our model; see Section 2.1.1) represents circulating mature blood cells. The intermediate compartments describe successive stages of progenitor cell differentiation.

In our framework, cells can only progress from one compartment to the next during reproduction. Fig 1 provides a schematic representation of the 27-compartment model for both healthy and cancer cells. Specifically, a cell in compartment *i* ∈ *{*2, …, 26*}* can undergo one of two events:

- *Differentiation*: the cell produces two daughter cells in the next compartment *i* + 1, occurring with probability *ϵ*_*H*_ for healthy cells and *ϵ*_*C*_ for cancer cells;
- *Symmetric division*: the cell produces two daughter cells that remain in compartment *i*, occurring with probability 1 − *ϵ*_*H*_ for healthy cells and 1 − *ϵ*_*C*_ for cancer cells.

**Fig 1.**
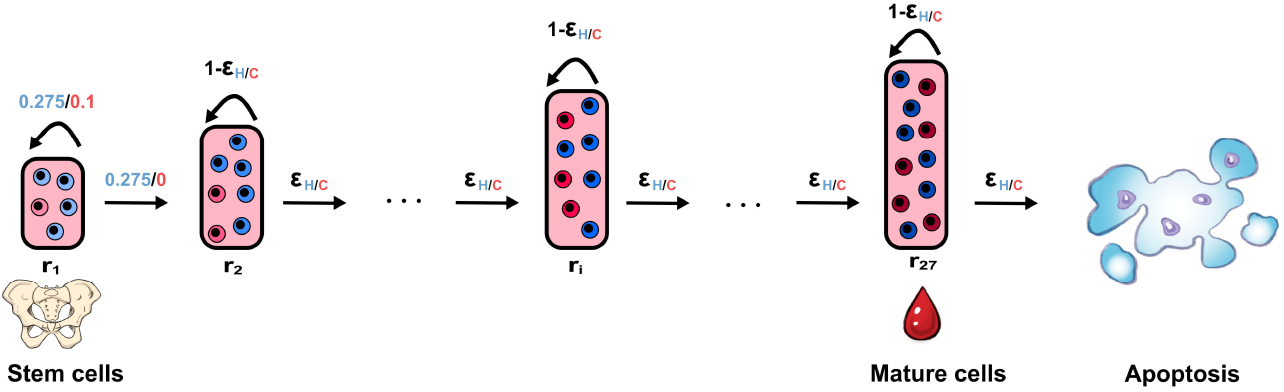
Schematic representation of the hierarchical 27-compartment model of CML. Healthy cells are represented in blue and cancer cells in red. Each compartment represents a stage of cellular differentiation. Cells undergo either symmetric division (probability 1 − *ϵ*_*H/C*_ ) or differentiation to the next compartment (probability *ϵ*_*H/C*_ ). Here, *ϵ*_*H*_ = 0.84 and *ϵ*_*C*_ = 0.67 denote differentiation probabilities for healthy and cancer cells, respectively. Cell numbers increase by a factor of 1.8 (rather than 2) between successive compartments to account for cell death in intermediate stages. A fictitious terminal compartment represents apoptosis (cell death in the blood). The replication rate *r*_*i*_ reflects the division speed in compartment *i*, ranging from *r*_1_ = 1*/*280 per day in stem cells to *r*_27_ = 5 per day in fully differentiated blood cells (see Eq (1)).

For compartment 27, we consider the same two events. Differentiation in this compartment leads to a terminal, fictitious compartment representing cell death in the bloodstream (apoptosis). The rate at which cells divide varies by compartment and is denoted by *r*_*i*_.

For stem cells (i.e. cells in the first compartment, *i* = 1), we assume that a third type of event can happen: *asymmetric cell division*. In this case, a cell in compartment *i* gives rise to one daughter cell that remains in the same compartment and another that moves to the next compartment (*i* + 1). The role of asymmetric division in the regulation and maintenance of the stem cell pool is discussed for instance in [22]. Asymmetric division is not thought to be a dominant mechanism in more differentiated blood cells, and thus it is not explicitly modelled in compartments beyond the first one.

However, a sequence of one symmetric division followed by one differentiation event can mimic the effect of asymmetric division, making it implicitly present in the model. Fig 2 provides a schematic representation of the three possible events in the first compartment for both healthy and cancer cells.

**Fig 2.**
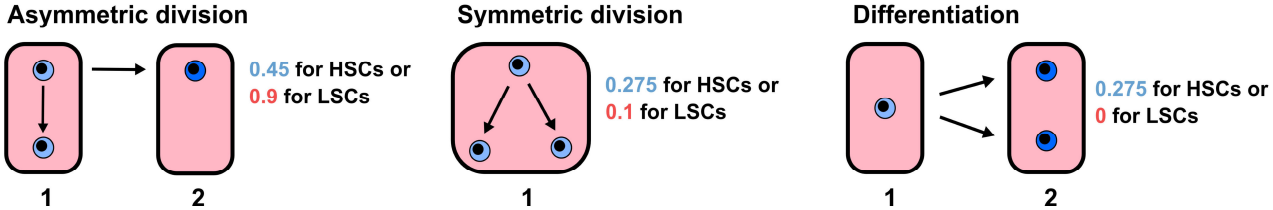
Schematic representation of the events in compartment 1. Hematopoietic stem cells are denoted by HSCs and leukemic stem cells by LSCs. There exist three types of events: asymmetric division (with probability 0.45 for HSCs and 0.9 for LSCs), symmetric division (with probability 0.275 for HSCs and 0.1 for LSCs), and differentiation (with probability 0.275 for HSCs and 0 for LSCs).

In the next two sections, we summarise the rationale behind the parameter values chosen for healthy and cancer cells. These values are detailed in a subsequent work in preparation.

#### 2.1.1 Parameters for hematopoiesis

Estimates of the number of hematopoietic stem cells (HSCs) in humans have been obtained through mathematical modelling. Lee-Six *et al*. [23] estimated the HSC pool to range between 50,000 and 200,000 cells. Observations from adult bone marrow indicate that over 70% of HSCs remain in a quiescent, non-dividing state [24]. This quiescence is thought to serve as a reserve mechanism, enabling rapid blood cell production in response to hematopoietic stress conditions such as severe infection, inflammation, or significant blood loss [25, 26]. Under normal physiological conditions, only the active (non-quiescent) fraction contributes to the continual renewal of mature blood cells. This is consistent with findings from both mouse and human studies suggesting that only a small number of cells would be required to recreate hematopoiesis after transplantation [27, 28]. Based on this evidence, we set the number of HSCs in healthy adults to 50,000 in our model (reflecting the active, non-quiescent fraction).

These HSCs divide on average once every 40 weeks, that is, approximately once every 280 days [29], and are responsible for maintaining an average of 2.8 *×* 10^11^ differentiated blood cells [30] per day. Building on the hematopoiesis framework proposed by Dingli *et al*. [14], we develop a 27-compartment stochastic model, in which each compartment represents a distinct stage of cell differentiation (see Fig 1 and Section 2.2). This number of compartments is of the same order of magnitude as what has been estimated for mice, which range from 15 to 22 differentiation stages [31], and is required to link the 50, 000 stem cells to the 2.8 *×* 10^11^ cells produced on average per day by the bone marrow.

Fig 3 illustrates how the 27 compartments in our model correspond to the main cell populations involved in hematopoiesis, namely, hematopoietic stem cells (HSCs), short-term HSCs (ST-HSCs), multipotent progenitors (MPPs), committed progenitors, precursors, and mature blood cells. The figure also shows the estimated number of cells per compartment on a logarithmic scale. To transition from approximately 50,000 stem cells to 2.8 *×* 10^11^ mature blood cells, 26 rounds of amplification are required. This quantitative expansion justifies the number of compartments chosen in our model.

**Fig 3.**
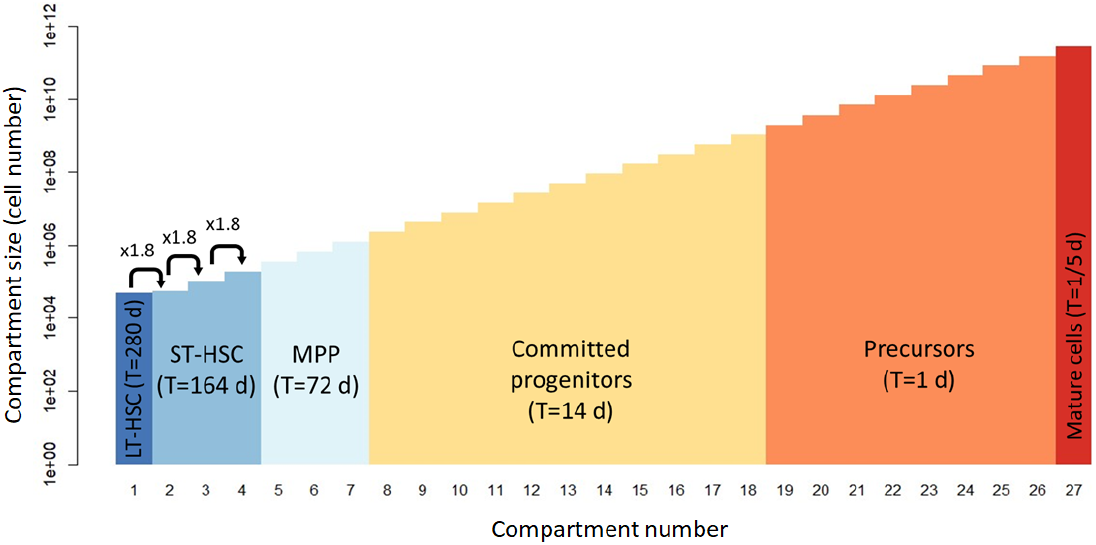
Logarithmic representation of the number of cells in each compartment. Compartment 1 contains 50,000 hematopoietic stem cells (HSCs), followed by compartments representing short-term HSCs (ST-HSCs), multipotent progenitors (MPPs), committed progenitors, and precursors. Compartment 27 corresponds to mature blood cells, with an estimated count of 2.8 × 10^11^. An amplification factor of *η* = 1.8 is observed between successive compartments. The time *T* between two successive divisions is shown in days (in brackets) for each population.

In the stem cell compartment, the probability of asymmetric division is estimated to be 0.45 [32]. Since the number of stem cells remains relatively constant over time, there must be as much symmetric division as differentiation in that compartment, which implies that the corresponding probabilities are both 0.275.

The probabilities of differentiation and symmetric division for healthy cells in compartment 2, …, 27, denoted by *ϵ*_*H*_ and 1 − *ϵ*_*H*_ respectively, are assumed to remain constant across all compartments of the hematopoietic system. For healthy cells, we use *ϵ*_*H*_ = 0.84, consistent with the value used in [14], despite differences in both the number of stem cells and the number of compartments.

To account for the varying pace of cell division — slower in HSCs and faster in mature blood cells — we introduce a compartment-specific replication rate *r*_*i*_, which represents the rate at which a cell undergoes either division or differentiation. This rate increases exponentially with the compartment index, following the relation *r*_*i*_ = *γ r*_*i*−1_ for *i* ≥ 2, with an estimated growth factor *γ* = 1.3213. This estimate of *γ* is close to the value 1.27 reported in [14]. Since HSCs divide roughly once every 280 days, we have *r*_1_ = 1*/*280, and the replication rate in compartment *i* is given by

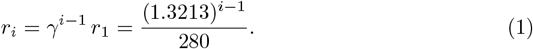

Because *γ >* 1, the replication rate increases with each compartment, resulting in increasingly larger compartments. The approximate size ratio between two successive compartments is 1.8, slightly less than 2, which reflects implicit cell loss due to death or differentiation failure. In addition, to prevent indefinite accumulation of cells in the final (27th) compartment, we explicitly incorporate a cell death process. In this compartment, each event results either in division (with probability 1 − *ϵ*_*H*_ ) or in apoptosis (with probability *ϵ*_*H*_ ), using the same value as for differentiation.

#### 2.1.2 Parameters for CML

In the remainder of the paper, we shift our focus to cancer cells. For these cells, all hematopoiesis parameters remain unchanged except for the following:

- The probabilities of asymmetric and symmetric cell divisions in the first compartment, estimated at 0.9 and 0.1, respectively, and the probability of differentiation, which is considered negligible.
- The probability of differentiation in compartments 2, …, 27, denoted by *ϵ*_*C*_, which is estimated at 0.67.

Since *ϵ*_*C*_ *< ϵ*_*H*_, there is reduced differentiation and therefore increased proliferation at various stages of cell maturation. This results in larger compartment sizes compared to the healthy case and accounts for the increased daily bone marrow production observed in CML patients: approximately 10^12^ cells per day [33], compared to 2.8 *×* 10^11^ in the healthy case [30].

### 2.2 Stochastic model for CML cell dynamics

In this section, we present the stochastic model we used to study cell dynamics across the 27 compartments of the hematopoietic system. This model is a specific type of branching process known as a *Markovian binary tree* (MBT), and plays a central role in investigating the disease initiation. We begin with a brief overview of MBTs before detailing their application to modelling the dynamics of both healthy and leukemic (CML) cells.

Branching processes are stochastic models that describe the evolution of populations in which individuals reproduce and die independently according to specified probability laws. These models have been widely applied in biology, ecology, epidemiology, and evolutionary theory, as well as in fields such as particle physics, chemistry, and computer science [34]. MBTs are a tractable class of continuous-time Markov branching processes, well-suited to modelling biological systems [35–37]. Originally introduced to describe evolutionary dynamics [38], MBTs track individuals whose lifetime and reproduction events are governed by an underlying continuous-time Markov chain with *n* ∈ ℕ transient states called *phases*, and one absorbing state. Phases may correspond to biologically meaningful attributes, such as age or health status, or serve as abstract intermediates. During their lifetime, individuals reproduce and move between phases until absorption (death). Birth and death rates typically depend on the current phase, and offspring start their life in a phase that may depend on the parent’s phase at the time of reproduction.

An MBT with *n* transient phases is governed by an underlying continuous-time Markov process characterised by three components: an *n × n* transition rate matrix *D*_0_, an *n × n*^2^ birth rate matrix *B*, and an *n ×* 1 death rate vector ***d***. Along with an initial probability vector ***α***, the matrices *D*_0_, *B*, and the vector ***d*** define the parameters of the model, and their entries have the following interpretations:

- *á*_*i*_ is the probability that the initial individual is in phase *i* at time *t* = 0;
- (*D*_0_)_*ij*_, for *i ≠ j*, is the rate at which an individual in phase *i* makes a transition to phase *j* without producing an offspring;
- *B*_*i,jk*_ is the rate at which an individual in phase *i* gives birth to a child and simultaneously changes to phase *j*, the child starting its life in phase *k*;
- *d*_*i*_ is the rate at which an individual in phase *i* dies (without producing offspring).

The diagonal entries of *D*_0_ are negative and such that *D*_0_**1** + *B***1** + ***d*** = **0**, where **1** denotes a column vector of ones of appropriate dimension. Fig 4 depicts a trajectory of an MBT initiated by a single individual. In practical applications of MBTs to real populations, the parameters ***α***, *D*_0_, *B* and ***d*** must be estimated from measurements obtained from the population.

**Fig 4.**
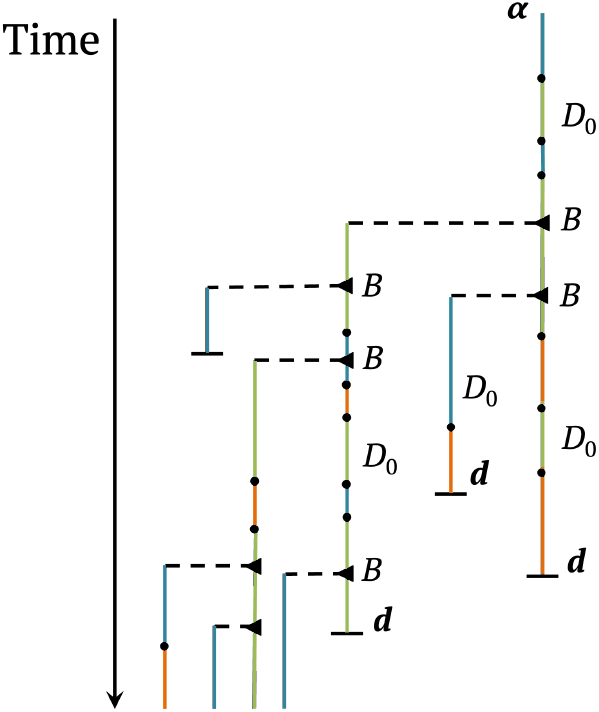
Schematic Illustration of the time evolution of a sample trajectory of a Markovian binary tree (MBT). This MBT depends on the parameters ***α***, *D*_0_, *B*, and ***d***. Vertical lines represent individual lifetimes, while dashed horizontal lines denote branching events (births). Coloured segments along the lifetime trajectories indicate transitions between transient phases of the underlying Markov process. Each individual eventually enters the absorbing phase and dies, indicated by a short terminal horizontal line. Given their initial phase, individuals evolve independently, with lifetime trajectories and reproduction governed by independent realisations of the same underlying Markov process

For *i* = 1, …, *n* and *t* ≥ 0, we let *Z*_*i*_(*t*) denote the number of individuals in phase *I* at time *t*, and we define the population size vector ***Z***(*t*) = (*Z*_1_(*t*), …, *Z*_*n*_(*t*)). The mean number of individuals in phase *j* at time *t*, conditional on starting from a single individual in phase *i*, is denoted by

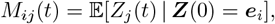

where ***e***_*i*_ is the *i*th unit vector of length *n*. The *n × n* mean population size matrix *M* (*t*) := (*M*_*ij*_(*t*)) takes the matrix exponential form

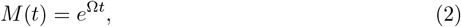

where the *n × n* matrix Ω is given by

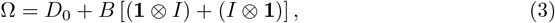

with **1** an *n ×* 1 vector of ones, *I* the *n × n* identity matrix, and ⊗ denoting the Kronecker product; we refer to [39] for derivations and further details. The unconditional mean population size vector ***m***(*t*) = (*m*_1_(*t*), …, *m*_*n*_(*t*)), where *m*_*j*_(*t*) := 𝔼 [*Z*_*j*_(*t*)], is given by

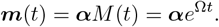

We use a specific MBT to model the progression of a population of CML cells throughout the 27 compartments of the hematopoietic system. Since cell replication rates depend on the compartment (see Section 2.1), we naturally adopt a modelling framework in which each compartment corresponds to a distinct phase of the MBT. Therefore, our model has *n* = 27 transient phases. In biology, a cell typically divides into two daughters. In the MBT framework, by contrast, it is often assumed that an individual gives birth to another while continuing its own life. For our purposes, the two views are equivalent, and we adopt the latter. Therefore, an ‘individual’ in our model, as represented by a vertical line in Fig 4, represents one cell lineage through successive compartments until cellular death from compartment 27.

The probability that the first leukemia cell arises in compartment *i* is given by the parameter *α*_*i*_. *The initial distribution vector* ***α*** *is unknown and is the quantity we aim to infer*. Based on the biological assumptions outlined in Section 2.1, the remaining model parameters are defined as follows:

- Since cells do not move between compartments without replicating, all off-diagonal entries of the transition rate matrix *D*_0_ are set to zero; that is, (*D*_0_)_*ij*_ = 0 for all *i ≠ j*.
- The non-zero entries of the birth rate matrix *B* reflect the possible outcomes of cell division, which depend on the current compartment. Cells may either divide symmetrically (producing two cells in the same compartment) or asymmetrically (one cell in the same compartment and one in the next). Differentiation occurs when both the daughter and the mother cells move to the next compartment. As a consequence, most entries of *B* are zero. Recalling that the replication rate in compartment *i* is given by *r*_*i*_ = *γ*^*i*−1^ *r*_1_, with *γ* = 1.3213 and *r*_1_ = 1*/*280, the non-zero entries are defined as follows:
- In the first row of *B*, corresponding to the stem cell compartment:

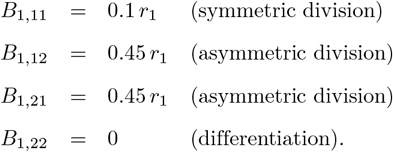

We have assumed that asymmetric division, which occurs with probability 0.9, is evenly split between the case where the daughter cell moves to compartment 2 and the mother remains in compartment 1, and the reverse. This symmetric splitting is an arbitrary modelling choice that does not affect the model outcomes, provided that *B*_1,12_ + *B*_1,21_ = 0.9 *r*_1_.

– In rows *i* = 2, …, 26, representing intermediate compartments:

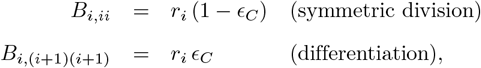

where *ϵ*_*C*_ = 0.67 is the differentiation probability for cancer cells.
– In the last row, corresponding to the 27th compartment:

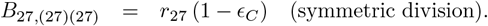

- Finally, we assume that cell death is negligible in all compartments except the last one. Thus, all entries of the death rate vector ***d*** are set to zero except for *d*_27_ = *γ*^26^ *r*_1_ *ϵ*_*C*_, which matches the differentiation rate in the final compartment.

With the phase-specific birth and death rates defined above, the diagonal entries of the transition rate matrix *D*_0_ are given by (*D*_0_)_*ii*_ = −*r*_*i*_; Since the model is Markovian, −(*D*_0_)_*ii*_ = *r*_*i*_ corresponds to the rate of the exponentially distributed time until a CML cell in compartment *i* undergoes a replication event (either division or differentiation).

Although this MBT is parametrised for a CML (leukemic) cell population, it can equally be used to model the dynamics of a healthy hematopoietic cell population. The structure of the model remains unchanged, the only differences lie in the value of the parameters, as shown in Fig 1 and Fig 2.

To efficiently simulate sample paths of the MBT, we use a hybrid stochastic-deterministic approach. Specifically, within each compartment, we simulate the population dynamics day by day using the Doob-Gillespie algorithm when the number of cells is below a threshold *V* = 150, since stochastic fluctuations are important when populations are small. Once the number of cells in a compartment exceeds this threshold at the end of a day, we approximate the dynamics by its expected value, computed from the mean population size matrix in Eq (2). This hybrid approach preserves randomness at low population sizes (when it is most important) and speeds up computations when populations become large.

Fig 5 and Fig 6 show simulated trajectories of compartment sizes in an MBT initiated with a single leukemia cell in compartment 1, 3, 5, or 10, which represent the disease progression in virtual patients. The stochastic fluctuations in the early compartments influence both the shape and magnitude of the population size curve corresponding to the final compartment 27. Since there is no differentiation in compartment 1 and a positive rate of symmetric division, the population in compartment 1 increases slowly on average. In compartments *i* = 2, …, 26 replication rates increase with the compartment index, and differentiation is more likely than symmetric division. As a result, each compartment increasingly ‘feeds’ the next one, leading to larger populations downstream. However, in the absence of sustained input from compartment 1, especially when the initial cell arises in a compartment later than 1, the population of cancer cells eventually dies out. This emptying effect is particularly visible in the trajectories of Fig 6, where no permanent self-sustaining growth occurs.

**Fig 5.**
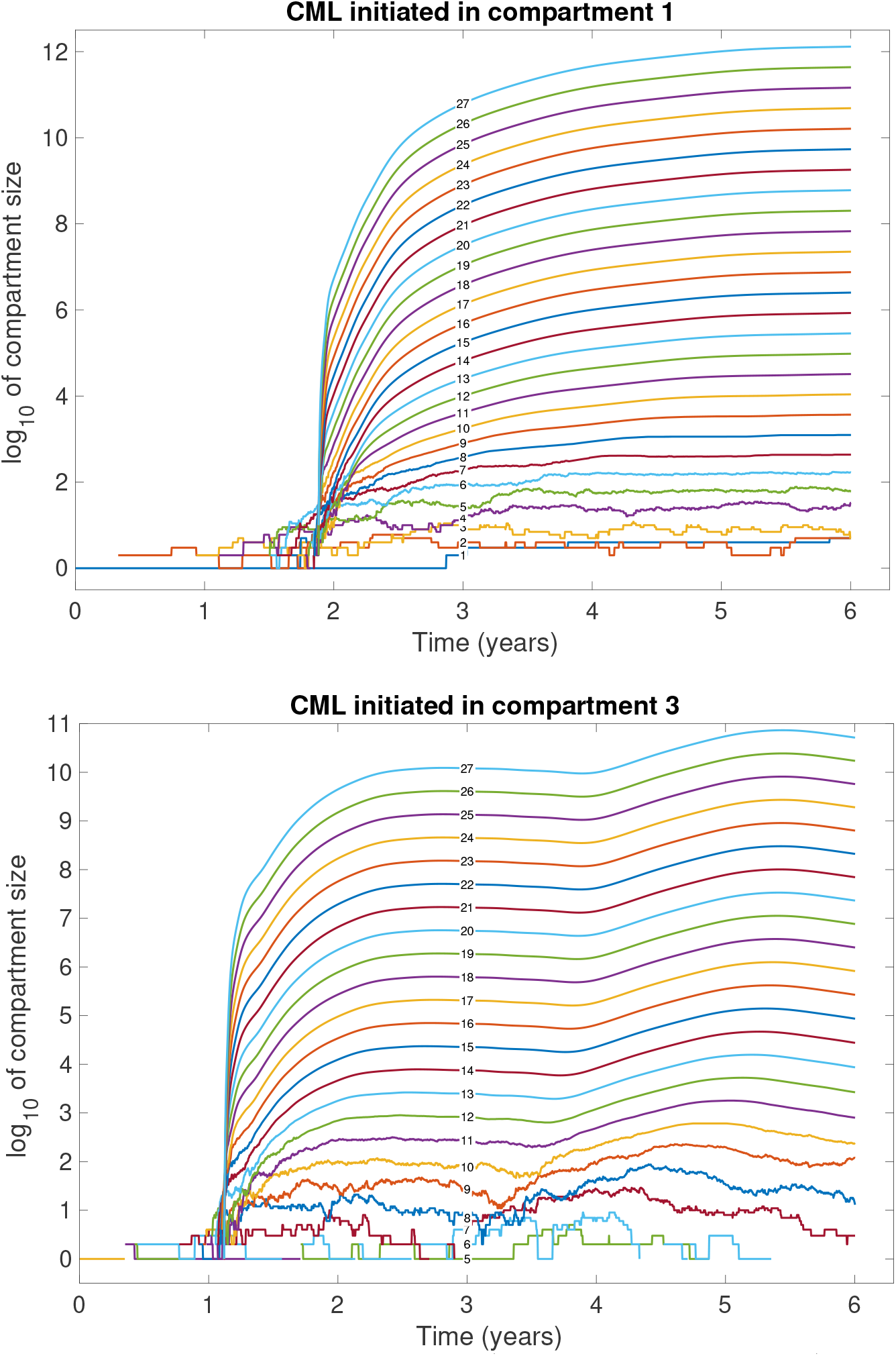
Simulated trajectories of compartment sizes (initiation in compartment 1 or 3). These simulated trajectories in an MBT initiated with one leukemia cell in compartment 1 (top) or 3 (bottom) represent the disease progression on virtual patients (in logarithmic scale). Trajectories that reach zero are omitted beyond that point, as log_10_(0) is undefined.

**Fig 6.**
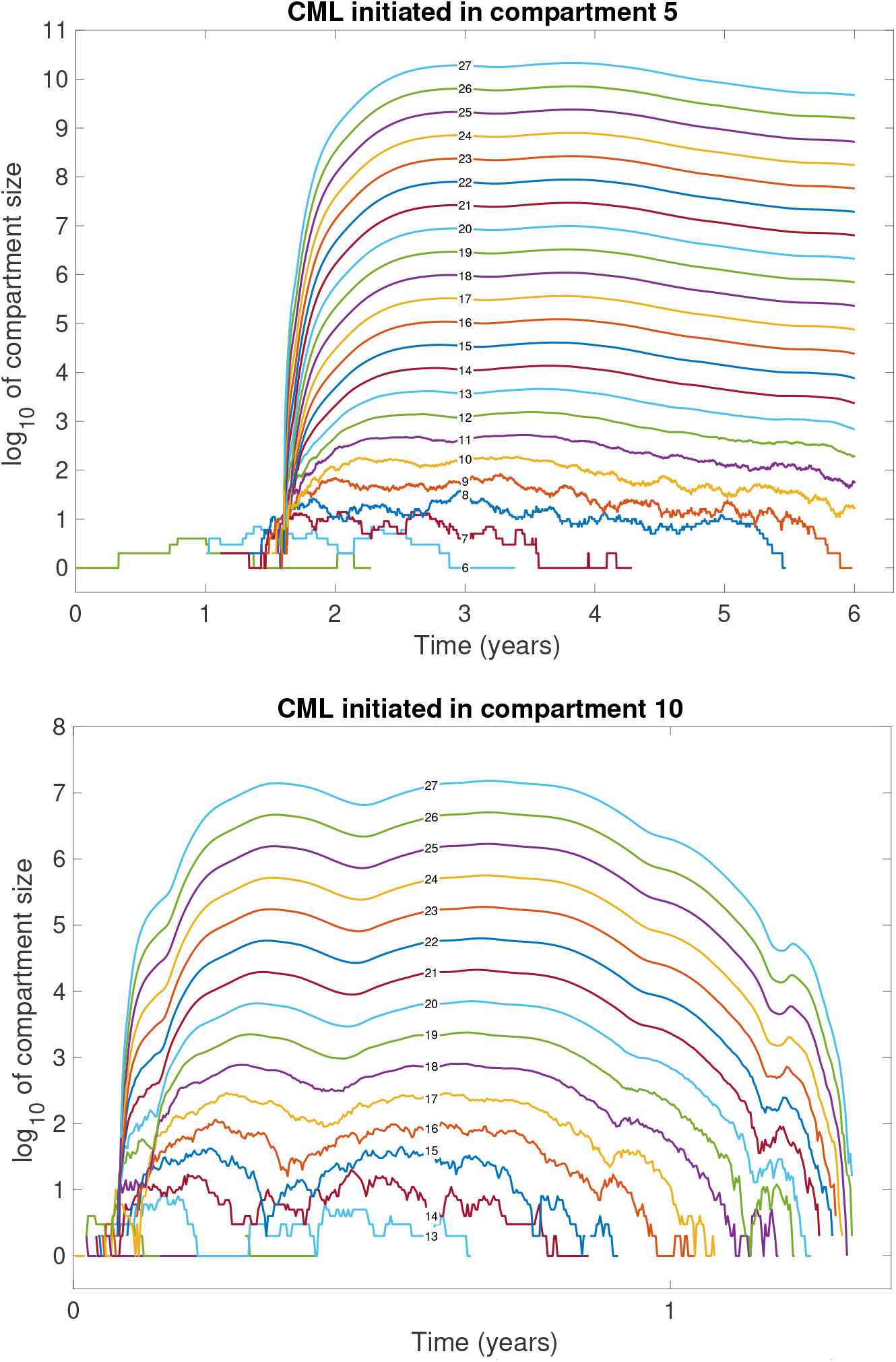
Simulated trajectories of compartment sizes (initiation in compartment 5 or 10). These simulated trajectories in an MBT initiated with one leukemia cell in compartment 5 (top) or 10 (bottom) represent the disease progression on virtual patients (in logarithmic scale). Trajectories that reach zero are omitted beyond that point, as log_10_(0) is undefined.

We also traced the mean curves using Eq (2) from our MBT model. The top panel of Fig 7 shows the results starting from a single cancer cell in compartment 1, while the bottom panel shows the results starting from a single cell in compartment 3. Our curves exhibit the same qualitative behaviour as those shown in Fig 4 of Werner *et al*. [16], which were obtained using a deterministic model. However, the numerical values differ slightly due to differences in parameter values.

**Fig 7.**
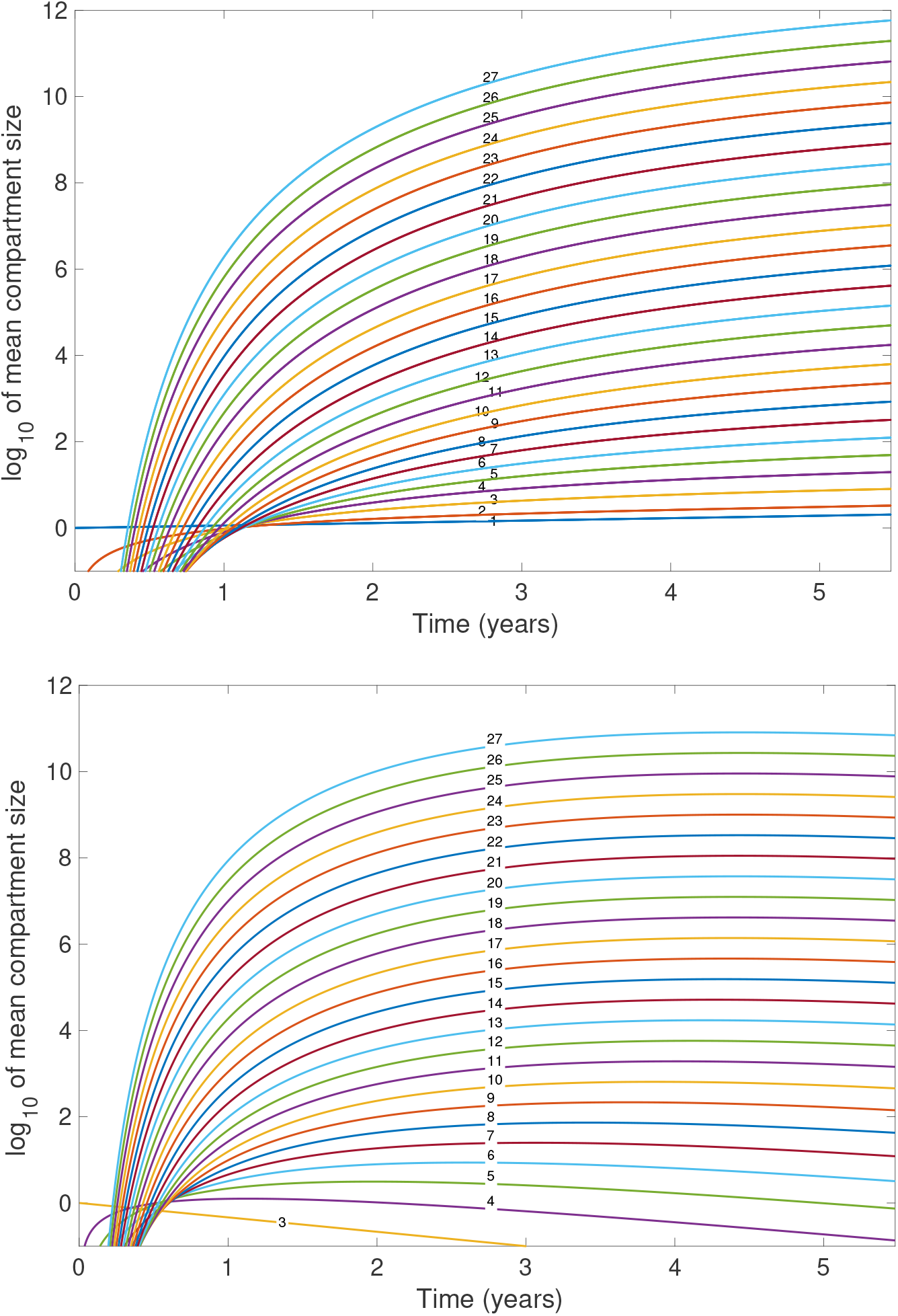
Mean number of cancer cells in each compartments (initiation in compartment 1 or 3). The mean curves represent the mean number of cancer cells in compartments 1 to 27 (in logarithmic scale) in an MBT initiated with one cancer cell in compartment 1 (i.e. one LSC; top) or with one cancer cell in compartment 3 (bottom).

## 3 Methods

### 3.1 Inference on the cellular origin of CML

CML most likely originates from a chromosomal rearrangement occurring in a single cell, as the probability of the same mutation arising independently in multiple cells is negligible [40]. In this section, we introduce a statistical method to infer the compartment in which the first CML cell emerged, based on observed counts of cancer cells in a patient’s blood at diagnosis. Our approach is based on a Bayesian framework known as *Approximate Bayesian Computation* with *Population Monte Carlo* (ABC–PMC).

To estimate the initial compartment distribution vector ***α***, we define a discrete parameter *θ* ∈ *{*1, 2, …, 27*}* which represents the compartment where CML initiates. We then interpret the posterior distribution of *θ* as an estimate for ***α***. Since ABC methods are traditionally used to estimate continuous parameters, we adapt the ABC–PMC algorithm to handle inference over bounded discrete parameter spaces.

We note that the estimated densities of cancer cell counts after six years obtained from the MBT model are multimodal and highly irregular (see Section 6.1), which makes likelihood-based inference of the disease origin unreliable. The ABC approach overcomes this limitation by directly estimating the posterior distribution over compartments, rather than relying on a single most likely origin.

In the remainder of this section, we start by presenting the standard ABC–PMC algorithm for continuous parameters, providing sufficient detail to help readers understand how the method works. We then describe how the method is adapted to accommodate inference for a discrete parameter.

#### 3.1.1 ABC - PMC for continuous parameters

Let *θ* be a model parameter taking values in a continuous space. The objective of Bayesian inference is to estimate the *posterior distribution f* (*θ* | *y*_obs_), which describes the distribution of *θ* given observed data *y*_obs_. By Bayes’ theorem, this posterior is given by

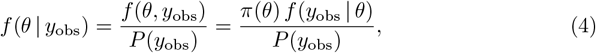

where *f* (*·* | *θ*) is the likelihood, *π*(*·*) is the *prior distribution*, and *P* (*·*) is the marginal likelihood (a normalising constant). Eq (4) represents the update of prior knowledge about *θ* after observing *y*_obs_. The posterior can also be written (up to a proportionality constant) as

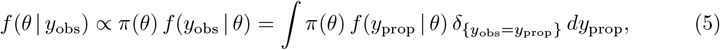

where *y*_prop_ denotes *proposed data*, that is, data simulated from the model using parameter value *θ*, and *δ* is the Dirac delta function.

*Approximate Bayesian Computation* (ABC) methods aim to approximate the posterior *f* (*θ* | *y*_obs_) when the likelihood is intractable or numerically costly to evaluate. ABC replaces the delta function in (5) by an approximate indicator. The resulting target distribution is

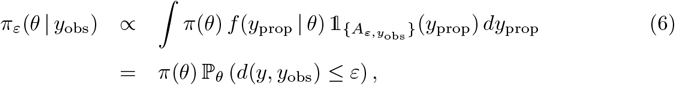

where 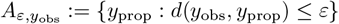 is a tolerance set, and *d*(*·, ·*) is a distance metric (for example, *d*(*y*_obs_, *y*) = |*y*_obs_ − *y*|). The indicator function 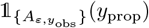 equals 1 when *y*_prop_ is sufficiently close to *y*_obs_ and 0 otherwise. ABC thus samples from an approximate posterior by retaining only those parameter values that generate simulated data sufficiently close to the observed data, within a tolerance *ε*.

Intuitively, the integrand in Eq (6) describes how accepted values of *θ* are obtained and contribute to the target distribution. First, we sample a value *θ*^*^ from the prior distribution *π*(*·*). Next, we generate proposed data *y*_prop_ from the model with parameter *θ*^*^, according to the likelihood *f* (*·* | *θ*^*^). Finally, we accept or reject parameter *θ*^*^ depending on whether the corresponding proposed data *y*_prop_ is ‘close enough’ to the observed data *y*_obs_. By repeating this procedure until we collect a total of *N* accepted values of *θ*, we obtain a population of parameter values, called *particles*, drawn from the target distribution. This procedure corresponds to the *ABC rejection method*.

The ABC rejection method can be computationally expensive for small values of *ε*, since fewer proposed samples are accepted as the tolerance threshold becomes narrower. Choosing an appropriate *ε* is crucial: smaller values improve the approximation of the true posterior but increase the computational cost.

To address this trade-off, a variant of the ABC rejection method can be used. It generates a sequence of intermediate distributions 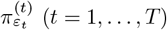, called *an ABC sequence*. This sequence gradually approaches the target distribution *π*_*ε*_(*·* | *y*_obs_) by using a decreasing sequence of tolerances *ε*_1_ *> · · · > ε*_*T*_. As a result, more proposals are accepted in earlier iterations (when *ε*_*t*_ is large). The approximation improves as *ε*_*t*_ decreases, but at the cost of increased computational effort. To balance the trade-off between accuracy and efficiency, the ABC sequence is often used in combination with *importance sampling* (described in the next paragraph), which helps reduce the number of samples needed to reach a desired level of accuracy. The resulting method is known as *ABC-Population Monte Carlo* (ABC–PMC).

In the ABC–PMC algorithm, at iteration *t*, we construct a posterior distribution 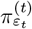 using the population of particles from the previous iteration. That is, we generate a new sample of parameter values 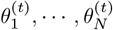 by building on the accepted particles from iteration *t* − 1, denoted 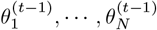, each associated with a weight 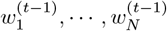 (these weights arise from importance sampling, as explained below). More specifically, at iteration *t*, we proceed as follows until we have accepted a new population 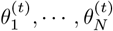 of *N* particles:

i. Select a parameter 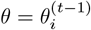 at random from the previous population, with probability proportional to its weight 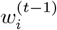;
ii. Perturb *θ* using a perturbation kernel *q*(*·* | *θ*), such as a normal distribution centred at *θ* (see below), to obtain a new candidate parameter *θ*^*^. Such a perturbation helps preserve particle diversity and allows the algorithm to explore the parameter space near good candidate solutions.
iii. Simulate proposed data *y*_prop_ from the model with parameter *θ*^*^. If the distance between *y*_prop_ and the observed data *y*_obs_ is less than the threshold *ε*_*t*_, then *θ*^*^ is accepted into the new population; otherwise, we return to step (i).

Using the law of total probability, the density of a new parameter value *θ*^(*t*)^ at iteration *t* (unconditionally on its acceptance) is given by

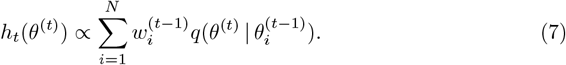

Note that, as we will see below, the weights depend on the choice of perturbation distribution. Because *h*_*t*_(*·*) differs from the prior distribution *π*(*·*), generating parameter values from *h*_*t*_ results in a biased posterior distribution:

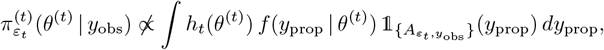

which does not match the target distribution in (6). To correct this bias, we use importance sampling and assign a new weight to each particle 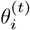 drawn from *h*_*t*_, defined as

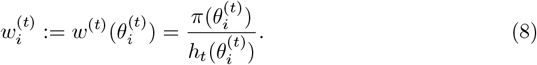

This correction ensures that the weighted sample approximates the target distribution:

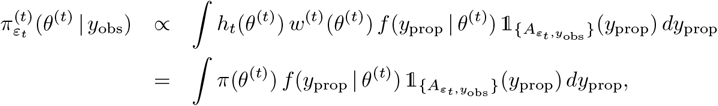

which now corresponds exactly to the target distribution (6).

The ABC–PMC algorithm is iterative, and several stopping criteria can be used: when the tolerance *ε*_*t*_ reaches a target value; when a predefined number *T* of iterations is completed; when the particle distribution stabilises; or when the acceptance rate drops below a given threshold (often 1%) [41].

#### 3.1.2 ABC–PMC for discrete parameters

We now assume that the parameter *θ* takes values in a finite discrete set 1, 2, …, *K*, and we describe how to adapt the ABC–PMC method to this setting. In our application to CML, the parameter *θ* corresponds to the compartment in which the first leukemic cell appears, with *K* = 27. The model used to generate proposed datasets *y*_prop_ is the MBT model introduced in Section 2.2. The observed data *y*_obs_ consist of leukemic cell counts in the blood of a patient at diagnosis, that is, the number of cancer cells present in compartment 27 six years after disease initiation. This six-year time frame is motivated by studies reporting a peak in leukemia cases in Hiroshima and Nagasaki in 1951, following the atomic bombings in 1945 [42, 43].

More recent findings suggest a wider and patient-specific latency period. In particular, Kamizela *et al*. [44] used phylogenetic reconstruction to estimate that the onset of the first malignant cell in CML typically occurs between 3 and 14 years prior to diagnosis. In Section 6, we explore the impact of this uncertainty by conducting a sensitivity analysis of the inferred initial CML compartment with respect to the assumed time from disease initiation to diagnosis.

In the discrete setting, the *N* particles sampled at each iteration of the ABC–PMC algorithm still represent parameter values, but these values now belong to the finite set 1, …, *K*. A key difference from the continuous case is that repeated values in the population are now possible, and even likely. For clarity, and to align with the CML application, we say that a particle “belongs to compartment *k*” if it takes the value *k*.

Here we can associate weights directly to compartments rather than individual particles. At each iteration *t*, we define the compartment weights as 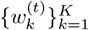, and the number of particles in compartment *k* as

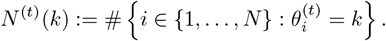

Since each particle must belong to one compartment, we have 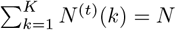. The posterior mass on compartment *k* is approximated by the normalised weight

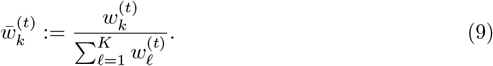

We assume a uniform prior *π*(*k*) = 1*/K* for *k* = 1, …, *K*. At iteration *t* = 1, we sample *N* particles 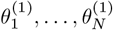 from the prior and retain those for which the model-generated data *y*_prop_ is sufficiently close to the observed data *y*_obs_. The initial (normalised) compartment weights are given by

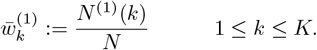

At each iteration *t* ≥ 2, we then proceed as follows:

i. Select a compartment *θ* = ℓ ∈ *{*1, …, *K}* at random with probability 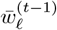.
ii. Perturb the selected compartment by drawing a new compartment *θ*^*^ = *k* from a discrete perturbation kernel *q*(*k* | ℓ) (see details below).
iii. Simulate data *y*_prop_ from the model with *θ*^*^ = *k*. Accept *θ*^*^ = *k* if *d*(*y*_prop_, *y*_obs_) ≤ *ε*_*t*_; otherwise, return to step (i).

Repeat this process until *N* accepted particles have been generated.

The marginal proposal distribution from which the values *θ*^(*t*)^ are drawn at iteration *t*, unconditionally on their acceptance, is given by

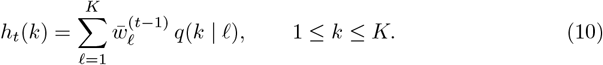

This is analogous to the continuous case, where particles are sampled from a mixture of perturbation kernels centred at previously accepted parameter values.

The importance weight for compartment *k* at iteration *t* is given by

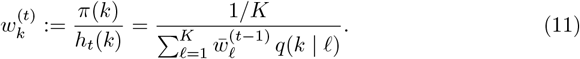

The weights are then normalised to obtain (9), which represents the estimated posterior mass associated with compartment *k* at iteration *t*. The collection of normalised weights 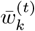 for *k* = 1, …, *K* provides a discrete approximation to the posterior distribution *π*_*ε*_(*θ* = *k* | *y*_obs_). A detailed justification for interpreting these normalised importance weights as posterior probabilities is provided in Section 6.2.

Thus, instead of assigning an individual weight 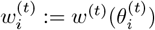 to each particle 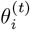 like in the continuous-parameter setting, here we assign a weight 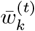 to each compartment *k*. The posterior distribution *π*_*ε*_(*·* | *y*_obs_) is then interpreted as a categorical distribution over the *K* compartments, and we obtain an estimator for the initial compartment distribution:

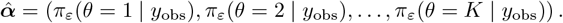

Each particle belonging to compartment *k* implicitly inherits the same weight 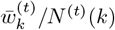. This discrete formulation offers an efficient and interpretable alternative to particle-wise weighting when the parameter space is finite and categorical. In practice, sampling and weighting can be performed either at the particle level or at the compartment level, with the two linked through *N* ^(*t*)^(*k*).

At iteration *t* ≥ 2, new particles are generated by resampling a previously accepted value *θ*^(*t*−1)^ and applying a perturbation kernel *q*(*k* | ℓ) to propose a candidate value *θ*^*^ in *{*1, …, *K}*. We consider two types of discrete perturbation kernels:

##### (i) Discretised normal proposal

To perturb a particle ℓ ∈ *{*1, …, *K}* in the discrete and bounded parameter space, we define a smooth proposal kernel based on a discretised normal distribution centered at ℓ with variance (*σ*^(*t*)^)^2^. For each candidate *k* ∈ *{*1, …, *K}*, the unnormalised transition probability is given by:

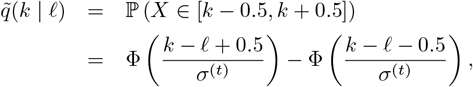

where *X* is a continuous normal random variable with mean ℓ and variance (*σ*^(*t*)^)^2^, and Φ(*z*) is the standard normal cumulative distribution function,

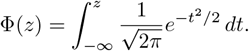

The kernel is then normalised over the support *{*1, …, *K}* to obtain a valid proposal distribution:

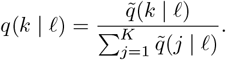

The result is a smooth, symmetric proposal that spreads mass across neighbouring values of ℓ, with the degree of spread controlled by *σ*^(*t*)^.

At iteration *t*, the variance (*σ*^(*t*)^)^2^ is adaptively updated as

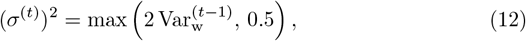

where 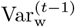 is the weighted variance of the previous particle population:

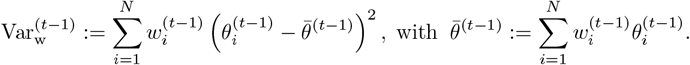

The definition in (12) ensures that the proposal remains diffuse enough to explore the parameter space (see for example [18]), while keeping the variance larger than 0.5 to avoid collapse in situations where all previous particles take the same value. In practice, since the parameter space is discrete and finite, it is often convenient to compute the weighted variance based on the posterior mass assigned to each compartment *k* ∈ 1, …, *K*:

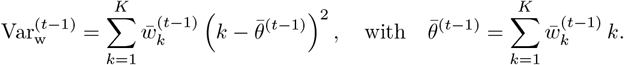

This formulation reduces the computational cost when the number of compartments *K* is much smaller than the number of particles *N*.

##### (ii) Symmetric random walk

We define a simple discrete random walk centered at the parent value ℓ which proposes a new value *θ*^*^ ∈ *{*ℓ − 1, ℓ, ℓ + 1*}* with equal probability. The corresponding proposal kernel is given by:

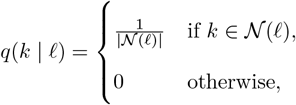

where the neighbourhood 𝒩(ℓ) := *{k* ∈ *{*1, …, *K}* : |*k* − ℓ| ≤ 1*}* adjusts for boundaries.

Algorithm 1 is an adaptation of the well-known ABC–PMC algorithm, tailored for the case of discrete parameters. In this setting, the problem can be interpreted as a model selection task, where each discrete value of the parameter *θ* corresponds to a distinct model. ABC approaches for model selection, such as the algorithm proposed by Toni *et al*. [45] (see also [46]), can be applied in this context. However, these methods are not sequential in nature and therefore differ from our proposed approach.

###### Algorithm 1

ABC–PMC for discrete parameter values with compartment weights

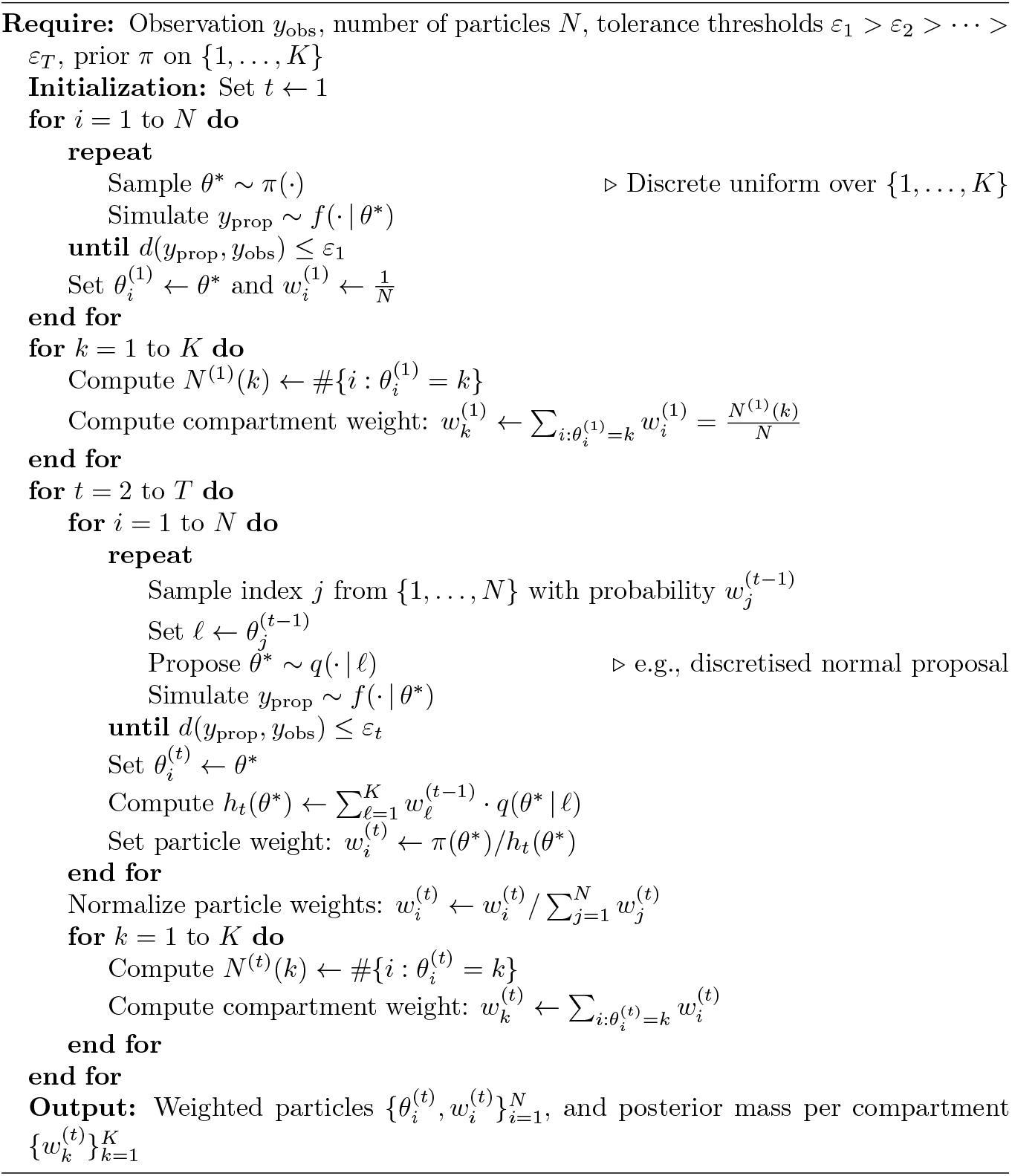

In the next section, we present numerical results for the ABC–PMC method for discrete parameters, and in Section 4.2, we apply it to infer the compartment in which CML likely originated in 19 patients from the TIDEL-II cohort [21].

## 4 Results

### 4.1 Validation using synthetic data

In this section, we apply the ABC–PMC method to identify the compartment (i.e., the stage of cell differentiation) at which the oncogenic *BCR::ABL1* protein first appeared, based on synthetic data.

A key strength of our model is that, at no additional cost, we can generate as many virtual patients as needed by simulating trajectories of the MBT model. Each trajectory represents a possible evolution of the disease, allowing us to efficiently explore a wide range of scenarios and outcomes.

To construct synthetic observations, we simulated four sets of 5000 MBT trajectories, each initiated with a single cancer cell in compartment *i*, for *i* = 1, 2, 3, or 4. From these, we extracted the median number of cancer cells in compartment 27 after six years for each of the four cases. These median values, which serve here as virtual observations in a proof-of-concept application of our ABC–PMC method, are respectively 6.7387 *×* 10^11^, 1.7357 *×* 10^11^, 1.4696 *×* 10^10^, and 2.0748 *×* 10^7^. For initial compartments greater than or equal to 5, the median is zero, since in over 50% of the trajectories, compartment 27 is empty after six years. Hence, we omit those cases from the analysis. We use medians rather than means because medians are more representative of typical model outcomes and are less sensitive to outliers, which are common in the highly skewed distributions produced by the stochastic simulations. To facilitate numerical stability and comparison, we work with the base-10 logarithm of the median cell counts. The corresponding log-scaled values are 11.8286, 11.2395, 10.1672, and 7.3170.

We set the number of particles to *N* = 200, the number of iterations to *T* = 15 (sufficient for the posterior distribution to stabilise), and use a threshold schedule defined by *ε*_1_ = 5 and *ε*_*t*_ = *ε*_*t*−1_*e*^−0.15^ for 2 ≤ *t* ≤ 15. The parameter space for the initial compartment *θ* is restricted to *{*1, 2, …, 8*}*. The results using the random walk perturbation kernel are shown in Fig 8 and Fig 9. The results obtained using the discretised normal perturbation kernel are similar. In all cases, we obtain a probability distribution over the possible compartments of origin, whose mode corresponds to the true origin of CML in our simulations.

**Fig 8.**
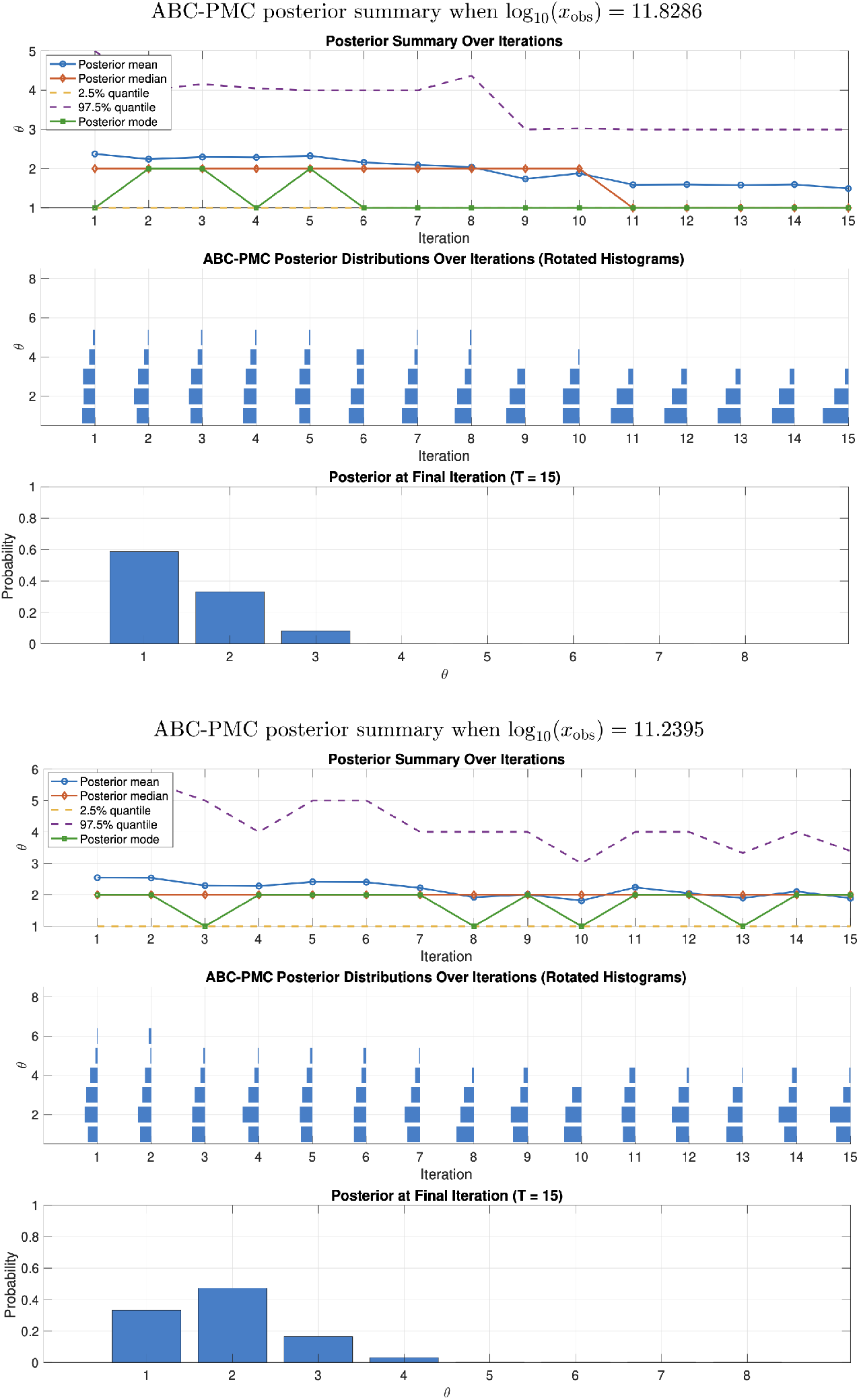
ABC–PMC posterior summary over iterations for the initial CML compartment (initiation in compartment 1 or 2). The observations correspond to the median number of cancer cells in compartment 27 after 6 years. Results are shown for simulations starting with one cancer cell in compartment 1 (top) or compartment 2 (bottom).

**Fig 9.**
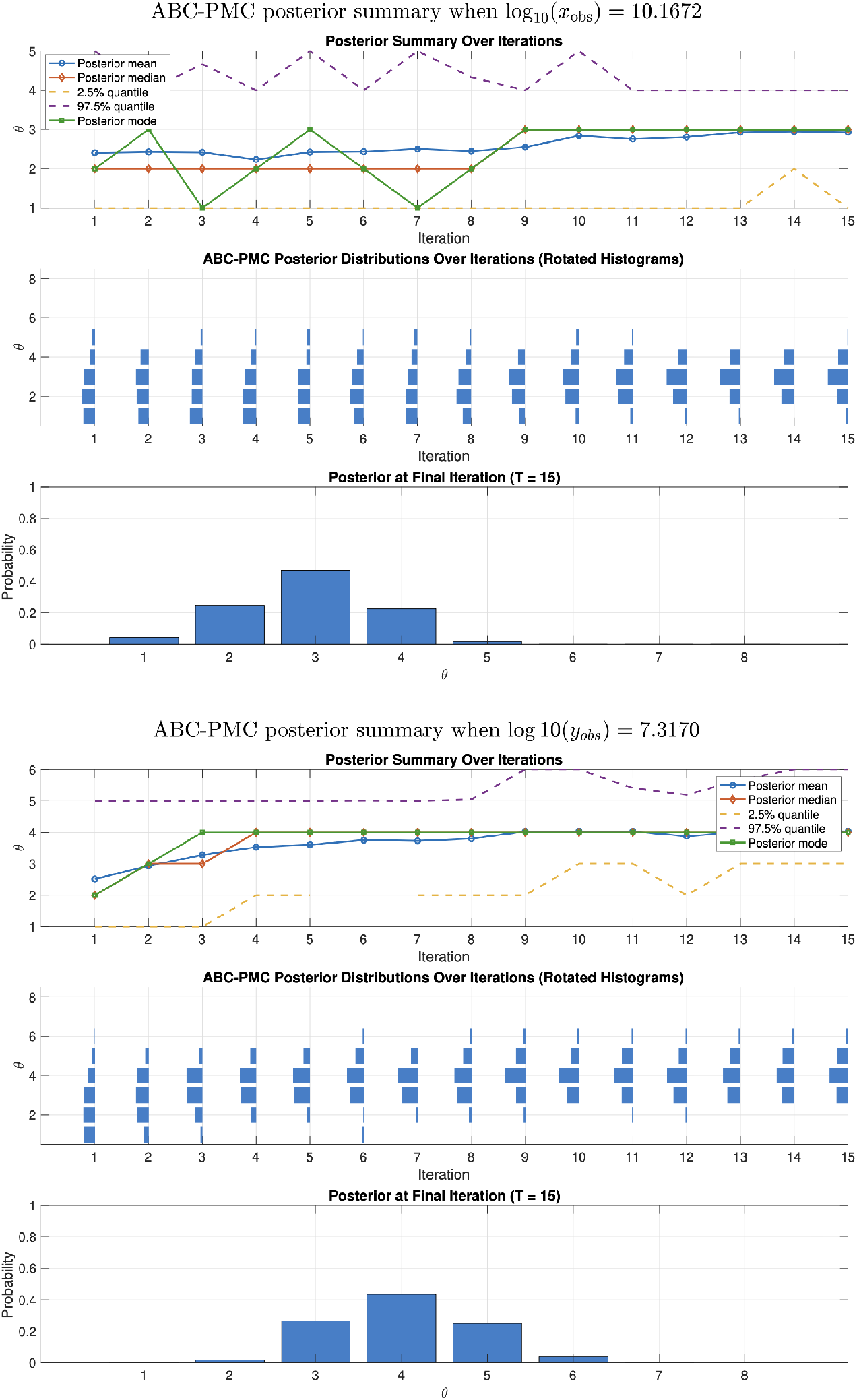
ABC–PMC posterior summary over iterations for the initial CML compartment (initiation in compartment 3 or 4). The observations correspond to the median number of cancer cells in compartment 27 after 6 years. Results are shown for simulations starting with one cancer cell in compartment 3 (top) or compartment 4 (bottom).

Having validated the method on synthetic data, we next apply it to clinical observations from the TIDEL-II cohort.

### 4.2 Application to patient data

In this section, we apply our methodology to a cohort of newly-diagnosed CML patients who responded to therapy from [21]. These data correspond to digital PCR (dPCR) performed on DNA taken from 19 patients. The advantage of quantification of *BCR::ABL1* DNA is that it is directly proportional to cell numbers, with one copy of *BCR::ABL1* in each cell. By quantifying *BCR::ABL1* and a control gene that is present both in CML and normal cells (on chromosome 7), glucuronidase beta (GUSB), the proportion of CML cells present in the blood (i.e. the last compartment in our model) at the time of diagnosis can be estimated, and the number *y*_obs_ of leukemic cells in the blood of a patient at diagnosis can be approximated. More precisely, the number of DNA copies for *BCR::ABL1* divided by the number of copies for GUSB (corrected by a factor of 2) provides information on the proportion of cancer cells relative to healthy cells for each patient. This percentage was then multiplied by 10^12^, the average number of total blood cells in patients with CML [47], to obtain the number of cancer cells per patient. In our model, we set the time of diagnosis at six years after the appearance of the first cancer cell, consistent with earlier studies of CML latency (see Section 3.1.2). We apply our ABC–PMC method for discrete values to these cancer cell estimates.

The results for one of the patients are shown in Fig 10 using the discretised normal perturbation kernel (top) and the random walk perturbation kernel (bottom). We set the number of iterations to *T* = 40, which provides sufficient stability in the posterior distributions. To reduce the noise induced by the stochastic simulations at each iteration, the reported posterior probabilities are computed as the average over the last 10 iterations rather than from the final iteration only.

**Fig 10.**
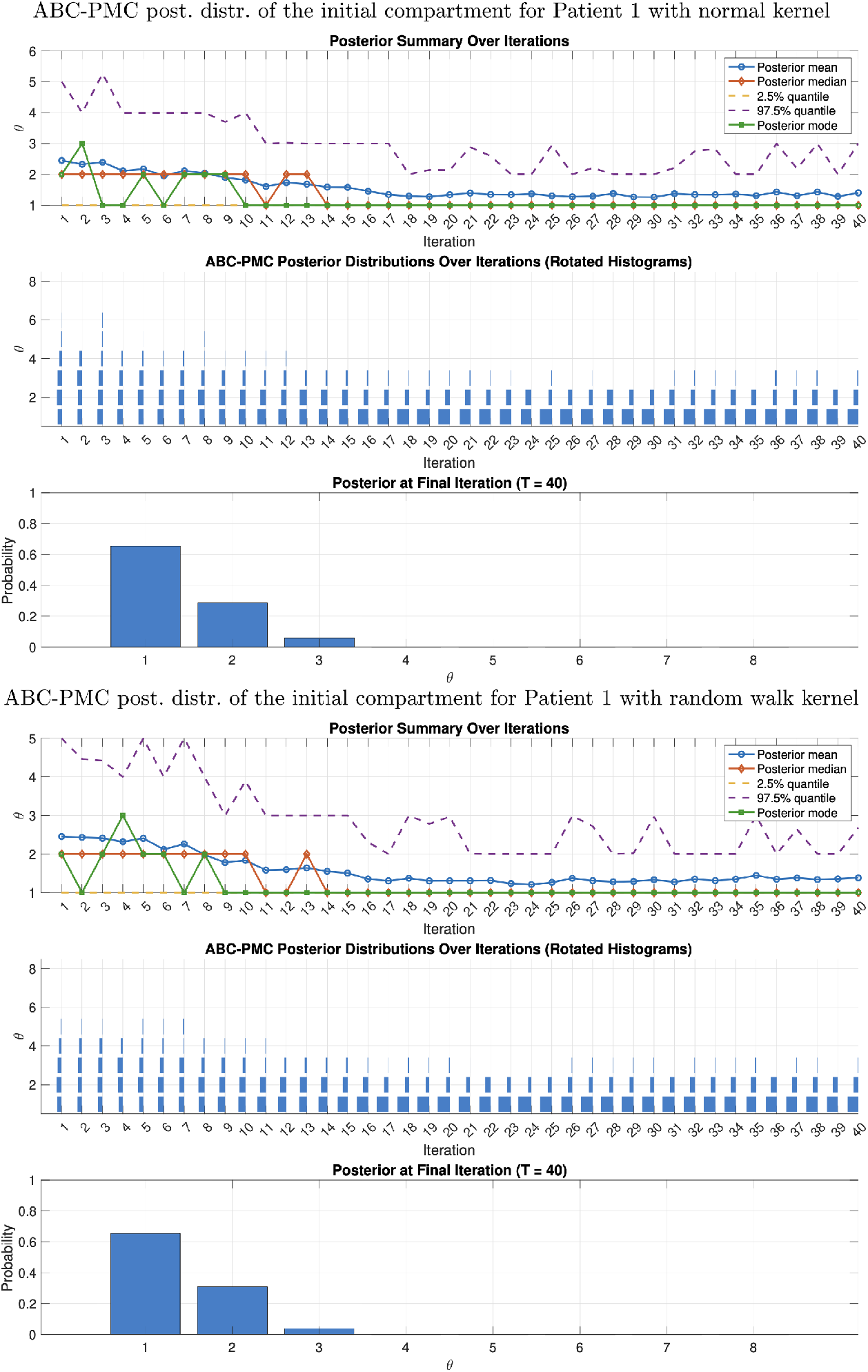
Posterior distribution for the origin compartment of CML for one patient. Results obtained for one of the patients of the TIDEL-II cohort [21] (Patient 1 in Table 1) with the discretised version of ABC–PMC with the discretised normal perturbation kernel (top) and the random walk perturbation kernel (bottom).

Both perturbation methods produce very similar posterior distributions: with the discretised normal kernel, the inferred (averaged) probability that the first cancer cell appeared in stem cell compartment 1 is 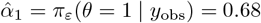, followed by 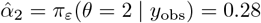 for compartment 2, and 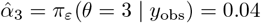 for compartment 3. With the random walk kernel, the corresponding probabilities are 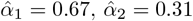, and 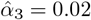. The posterior probabilities obtained using the discretised normal kernel are reported for all 19 patients in Table 1.

**Table 1.** Observed number of cancer cells *y*_obs_ (in units of 10^11^ and in logarithmic form) for 19 patients from the TIDEL II cohort [21], together with the posterior probabilities 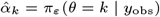 of CML initiation in compartment *k* (for *k* = 1, 2, 3) for each patient. These probabilities were obtained using the ABC–PMC algorithm with the discretised normal perturbation kernel for *T* = 40 iterations. Reported values correspond to the average over the last 10 iterations to reduce Monte Carlo noise.

| **Patient** | $y_{\text{obs}}/10^{11}$ | $\log_{10}(y_{\text{obs}})$ | $\hat{\alpha}_1$ | $\hat{\alpha}_2$ | $\hat{\alpha}_3$ |
| --- | --- | --- | --- | --- | --- |
| 1 | 6.84 | 11.8352 | 0.68 | 0.28 | 0.04 |
| 2 | 4.45 | 11.6488 | 0.56 | 0.37 | 0.07 |
| 3 | 4.87 | 11.6879 | 0.59 | 0.34 | 0.07 |
| 4 | 6.58 | 11.8181 | 0.68 | 0.27 | 0.05 |
| 5 | 5.17 | 11.7134 | 0.66 | 0.30 | 0.04 |
| 6 | 6.64 | 11.9367 | 0.84 | 0.16 | 0 |
| 7 | 5.68 | 11.7542 | 0.70 | 0.28 | 0.02 |
| 8 | 4.78 | 11.6792 | 0.59 | 0.36 | 0.05 |
| 9 | 8.88 | 11.9486 | 0.90 | 0.10 | 0 |
| 10 | 11.0 | 12.0422 | 0.90 | 0.09 | 0.01 |
| 11 | 12.4 | 12.0934 | 0.86 | 0.14 | 0 |
| 12 | 5.99 | 11.7774 | 0.69 | 0.27 | 0.04 |
| 13 | 6.30 | 11.7993 | 0.70 | 0.27 | 0.03 |
| 14 | 6.80 | 11.8325 | 0.70 | 0.28 | 0.02 |
| 15 | 6.47 | 11.8109 | 0.69 | 0.27 | 0.04 |
| 16 | 16.4 | 12.2146 | 0.96 | 0.04 | 0 |
| 17 | 12.1 | 12.0824 | 0.87 | 0.12 | 0.01 |
| 18 | 9.38 | 11.9722 | 0.89 | 0.11 | 0 |
| 19 | 9.77 | 11.9899 | 0.89 | 0.10 | 0.01 |

We can make several observations from Table 1. First, across the entire cohort, compartment 1 (stem cells) consistently appears as the most likely origin of CML. However, the non-zero values of 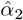 and 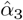 highlight the fact that we cannot exclude the possibility that CML is initiated in compartments 2 or 3, and they quantify the likelihood of these alternative origins for each patient.

Second, over the range of observations for the 19 patients from the TIDEL II cohort (i.e. for log_10_(*y*_obs_) between 11.6488 and 12.2146), linear relationships appear to emerge between the logarithm of the number of cancer cells at diagnosis, log_10_(*y*_obs_), and the estimated probabilities 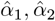, and 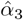 that the disease originated in compartments 1, 2 or 3.

To test this relationship further, we extended the range and added virtual observations (not corresponding to real patients). We obtained strong evidence of a linear relationship over the range of patient data, yielding the following explicit estimators for the probabilities that the first cancer cell originated in compartment *k*, for *k* = 1, 2, 3 (constrained to take values in [0, 1]):

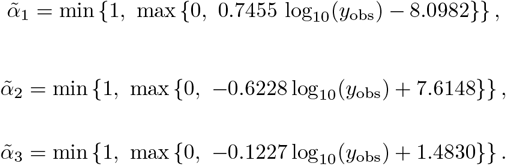

Outside the range of patient data, the relationship generally extends to a logistic form for 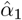 and 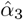, while the relationship for 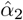 is less clear. Full details and figures are provided in Section 6.3.

The results in Table 1 were obtained under the assumption that diagnosis occurs six years after disease initiation. More recent studies suggest a broader window of 3–14 years before diagnosis [44]. In Fig 11, we assess the sensitivity of our results to the assumed timing of diagnosis. We focus on four representative patients: Patients 2 and 8 (with relatively low 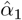 and high 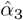), Patient 14 (with an average profile), and Patient 16 (with relatively high 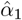). We report posterior probabilities for diagnosis occurring between four and twelve years after disease initiation, noting that the general trends can be extrapolated beyond this range. We observe that the stem cell compartment 1 remains the most likely origin of the disease in all cases. For Patient 2, however, the probability that CML originated in compartment 2 approaches that of compartment 1 when the diagnosis is assumed to occur between eight and ten years after disease initiation.

**Fig 11.**
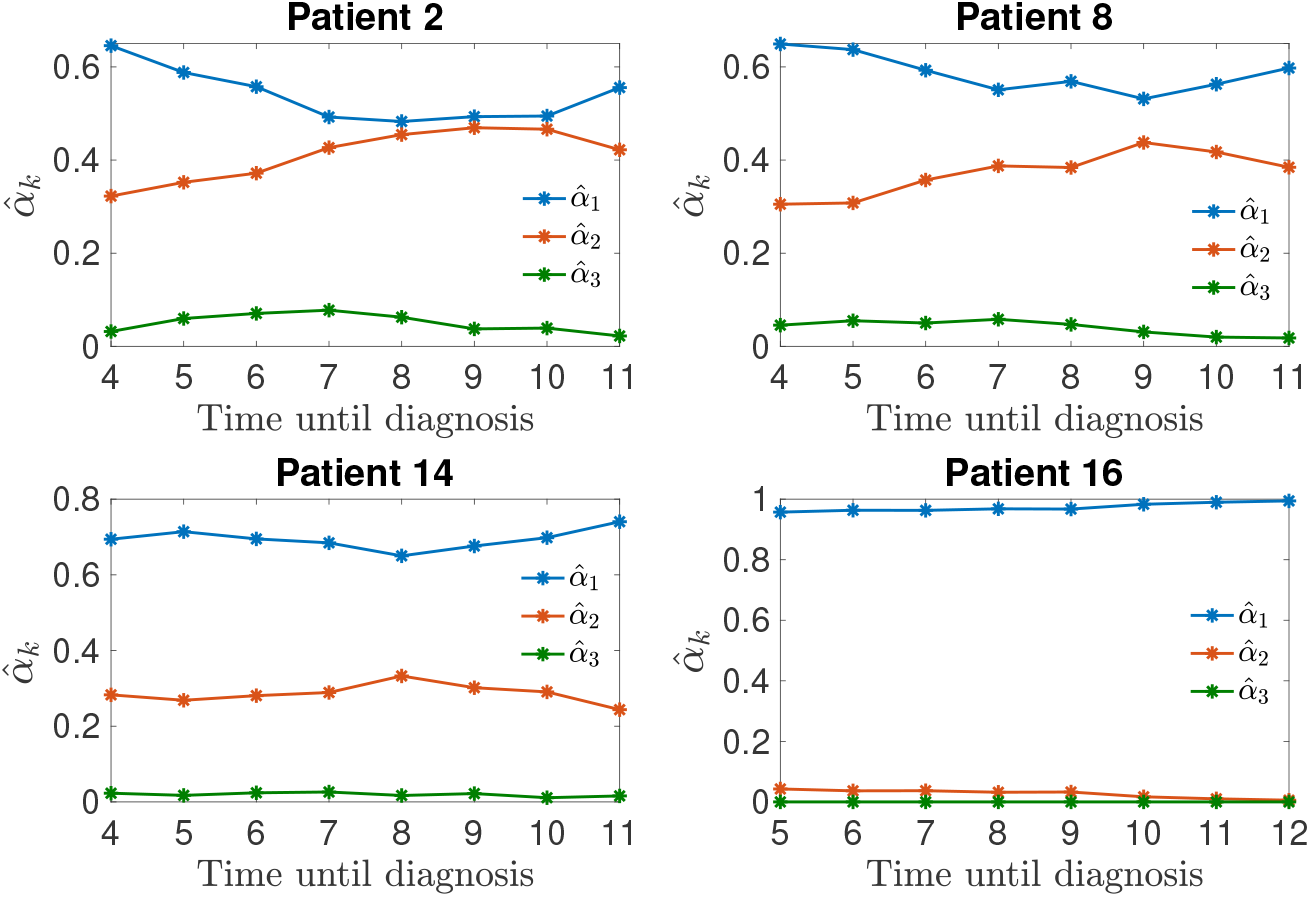
Posterior probabilities of CML initiation for different diagnosis times (in years). Posterior probabilities 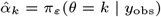 of CML initiation in compartment *k* (*k* = 1, 2, 3) for selected patients, obtained using the ABC–PMC algorithm with the discretised normal perturbation kernel over *T* = 40 iterations. Results are shown for a range of assumed times until diagnosis. Values correspond to the average over the last 10 iterations to reduce Monte Carlo noise.

*Remark:* In these experiments, we used an adaptive version of the ABC–PMC algorithm, where the tolerance at each iteration was updated using the previous value and a chosen quantile of the simulated distances, allowing the tolerance to decrease gradually while remaining sufficiently large when the model was less well aligned with the data (e.g., for diagnosis times differing substantially from six years).

Our results suggest that CML most likely originates in hematopoietic stem cells (HSCs), or in cells that closely resemble stem cells in their ability to divide and differentiate. An important open question is whether the first leukemic cells are truly stem cells, or if they are more differentiated progenitor cells that have somehow reverted to a stem-like state. Experimental evidence on this point is mixed: one study demonstrated that committed progenitor cells (specifically, granulocyte-macrophage progenitors) can be reprogrammed into leukemia stem cells (LSCs) by introducing a specific fusion gene, MLL-AF9 [48, 49]. However, another study found that the oncogene *BCR::ABL1* (the driver mutation in CML) cannot trigger this kind of reprogramming in committed progenitors [50]. Since additional mutations beyond *BCR::ABL1* are found at diagnosis in a minority of CML patients [51, 52], it is more likely that the cancer originates directly in normal stem cells that have been transformed. This interpretation aligns with our inference that the disease typically begins in the stem cell compartment.

Moreover, the presence of leukemic cells in the stem cell compartment is consistent with experimental evidence showing that the *BCR::ABL1* fusion gene is expressed not only in myeloid cells, but also in cells of the lymphoid lineage, such as T and B lymphocytes [53]. This suggests that the oncogenic event occurs early in hematopoiesis, before lineage commitment. In addition, LSCs have been detected in the bone marrow of all CML patients studied [54], further supporting a stem-cell origin.

In summary, by combining a stochastic model of CML progression (MBT) with a Bayesian inference framework (ABC–PMC), we have addressed the question of disease origin. Our results strongly indicate that CML most likely arises in the stem cell compartment. Taken together with the biological literature, this suggests that LSCs likely derive directly from healthy hematopoietic stem cells (HSCs), rather than from more differentiated progenitors.

## 5 Discussion

Previous studies have shown that the development of both normal and leukemic blood cells can be represented using a multicompartment model, where each compartment corresponds to a stage of cellular differentiation, from hematopoietic stem cells (HSCs) to mature blood cells [14, 16]. Building on this approach, we used a class of stochastic branching processes known as Markovian binary trees (MBTs) [55] to model the dynamics of healthy and leukemic cell populations across 27 compartments. Despite its relatively simple parameterisation, the MBT model captures key features of hematopoiesis, including the exponential growth of leukocytes and the disruption of normal differentiation observed in leukemia.

Although much is known about the biology of chronic myeloid leukemia (CML), the precise cellular origin of the leukemia-initiating cell, or leukemia stem cell (LSC), remains debated [17]. While the term “LSC” implies a stem cell origin, some mouse models suggest that committed progenitors can acquire stem-like self-renewal capacity following genetic reprogramming events [17, 48]. Conversely, other studies have shown that the *BCR::ABL1* oncogene — the primary molecular driver of CML — can provide a proliferative advantage, but is insufficient to induce self-renewal in committed progenitors [50].

To address this question, we developed a discrete-parameter version of the Approximate Bayesian Computation with Population Monte Carlo (ABC–PMC) algorithm [19]. In our framework, the discrete parameter represents the compartment (i.e., stage of differentiation) in which the first leukemic cell appeared, and the observed data consist of leukemic cell counts measured in the most mature compartment (compartment 27) at the time of diagnosis, as inferred from dPCR assays on patient blood samples.

We first validated our approach using synthetic data and then applied it to clinical data from 19 newly-diagnosed patients [21]. The posterior distributions consistently support the hypothesis that the initial leukemic cell arose in the stem cell compartment or in a cell exhibiting stem-like properties. We assumed a time-to-diagnosis of six years following disease initiation, an estimate supported by epidemiological data from Hiroshima and Nagasaki survivors [42, 43]. Sensitivity analyses demonstrated that small deviations in this assumption do not substantially alter the inference for early compartments, confirming the robustness of our conclusions. Interestingly, we cannot exclude the possibility that in some cases the initial leukemic cell arose in compartment 2 or 3, leading to a longer latency before diagnosis. A recent study using mutational data to reconstruct the phylogeny of CML inferred that latency is more variable than commonly assumed [44]. Our model indicates that the cell compartment of origin is one possible explanation for variable growth rates.

By modelling the full course of the disease, this work opens the way for addressing new questions, notably regarding optimal treatment duration and relapse risk following treatment discontinuation. These questions will be addressed in a subsequent work (manuscript in preparation).

The discrete ABC–PMC framework we have developed can be extended to other settings. One promising direction is the study of pre-treatment resistance in CML [56–58], which would require estimates of the number and timing of resistant clones at diagnosis, which could potentially be derived from dPCR and clonal mutation data. Existing phylogenetic approaches use allele frequency data to reconstruct mutational timelines [23, 59, 60]; our method could complement these models by estimating the timing or location of key mutational events in a probabilistic framework. More broadly, the ABC–PMC method for discrete parameters can extend beyond CML and find applications in various scenarios involving discrete parameters.

## 6 Supporting information

### 3.1 Approximate densities of cancer cell counts under different initiation compartments

Fig 12 shows the empirical distributions of the number of cancer cells in compartment 27 after six years, obtained from 10,000 simulated trajectories of the MBT model. The top panel corresponds to simulations starting with one cancer cell in compartment 1 (stem cells), and the bottom panel to simulations starting with one cancer cell in compartment 2. The estimated densities are highly irregular and multimodal, reflecting the stochastic variability in cell population growth. These non-smooth, multi-spike patterns illustrate why likelihood-based estimation of the compartment of origin is unreliable and motivate the use of the ABC framework, which does not rely on explicit likelihood evaluation.

**Fig 12.**
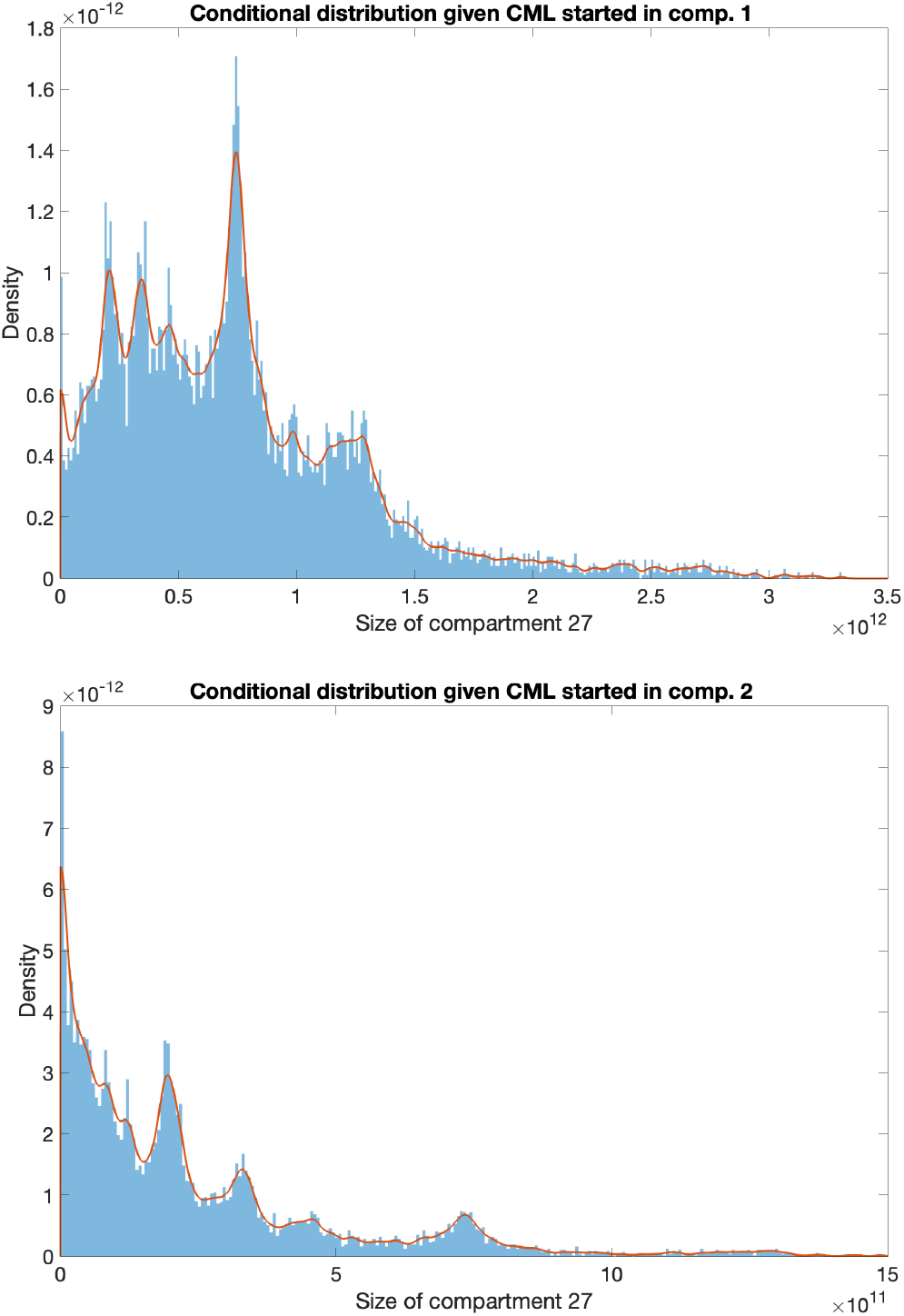
Approximate densities of cancer cell counts in compartment 27 after six years. These densities were obtained from 10,000 MBT simulations starting from compartment 1 (top) and compartment 2 (bottom). The multimodal, irregular shape of these distributions highlights the difficulty of applying likelihood-based inference and motivates the ABC approach.

### 6.2 Importance sampling weights in ABC–PMC for discrete parameters

Let *θ* be a discrete parameter taking values in *{*1, 2, …, *K}*. In the ABC framework, we aim to approximate the ABC posterior distribution

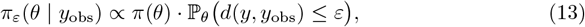

where *π*(*θ*) is the prior distribution, *y* ∼ *f* (*·* | *θ*) is data simulated from the model, *d*(*·, ·*) is a distance function, and *ε >* 0 is a tolerance threshold (see Eq (6)). When the likelihood *f* (*·* | *θ*) is intractable or computationally expensive to evaluate, ABC avoids evaluating it directly. Instead, it uses simulations from the model to estimate how likely it is to observe data similar to the actual data *y*_obs_, as measured by the criterion *d*(*y, y*_obs_) ≤ *ε*. This leads to a likelihood-free approximation of the posterior that depends only on the prior and the simulated distance probabilities.

In the ABC–PMC algorithm, the approximation is refined iteratively. At each iteration *t*, a set of *N* parameter values *θ*_1_, …, *θ*_*N*_ is sampled from a proposal distribution *h*_*t*_(*θ*), typically constructed using the particles accepted at the previous iteration. For each sampled *θ*_*i*_, a simulated dataset *y*_*i*_ is generated and *θ*_*i*_ is accepted only if the distance *d*(*y*_*i*_, *y*_obs_) ≤ *ε*_*t*_. Thus, the *accepted* samples are drawn from the proposal distribution *conditioned on acceptance*:

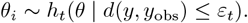

The accepted *θ*_*i*_ are effectively distributed according to the proposal distribution re-weighted by the ABC acceptance probability:

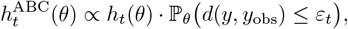

while our target distribution is the ABC posterior (13) with *ε* = *ε*_*t*_ at iteration *t*:

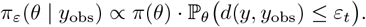

To adjust for the difference between the target and the proposal, we apply importance sampling [61–63]. The resulting unnormalised importance weight 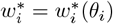 of each accepted *θ*_*i*_ is obtained by considering

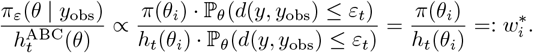

This shows that, although the simulated likelihood does not appear explicitly in the weights, its influence is already accounted for through the ABC acceptance step. The unnormalised weights can then be normalised:

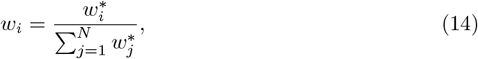

and used to approximate expectations under the ABC posterior using the standard *importance sampling identity*. For any function *f* (*θ*), the expectation of *f* (*θ*) under the ABC posterior can be written as

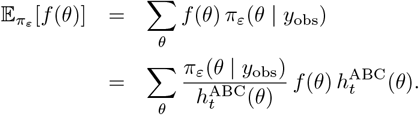

This can be approximated via Monte Carlo using the *accepted samples*:

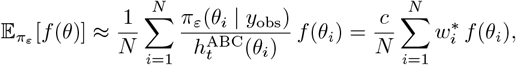

for a normalising constant *c*, since the weights 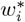 are unnormalised. Taking *f* (*θ*) = 1 yields 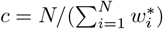. This means that, using the normalised weights (14),

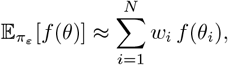

which removes the explicit factor 1*/N* and is more common in ABC implementations (see for instance [61, Section 3.3.2]).

To estimate the ABC posterior mass at a particular discrete value *k* ∈ 1, …, *K*, we use the indicator function *f* (*θ*) = **1**_*θ*=*k*_. Then:

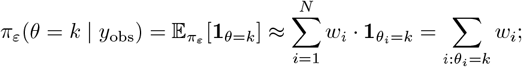

(see for example [63, Section 2.2] for the continuous case). Hence, the posterior mass at *θ* = *k* is approximated by summing the normalised weights of the particles with value *θ*_*i*_ = *k*. This identity is a direct consequence of the importance sampling approximation and justifies interpreting the weights as a discrete approximation to the ABC posterior distribution over *θ*.

### 6.3 Linear and logistic relationships between 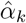 **and** log_10_(*y*_**obs**_)

Fig 13 depicts, with pink disks, the estimated values of 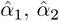, and 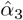 obtained using the ABC–PMC algorithm, plotted as functions of the logarithm of the number of cancer cells at diagnosis, log_10_(*y*_obs_), for the 19 TIDEL-II patients from [21].

Over the range of the patients data, we found a strong and statistically significant positive linear relationship between log_10_(*y*_obs_) and 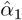, with a Pearson correlation coefficient of *r* = 0.9456 and a p-value of *p* = 1.0249 *×* 10^−9^. In contrast, 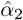 and 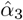 showed strong and statistically significant negative linear relationships (*r* = −0.9504, *p* = 4.73 *×* 10^−10^ for 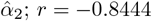, *p* = 5.46 *×* 10^−6^ for 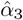), indicating clear inverse associations. The red plain lines in Fig 13 correspond to the linear regression fits reported in Section 4.2.

**Fig 13.**
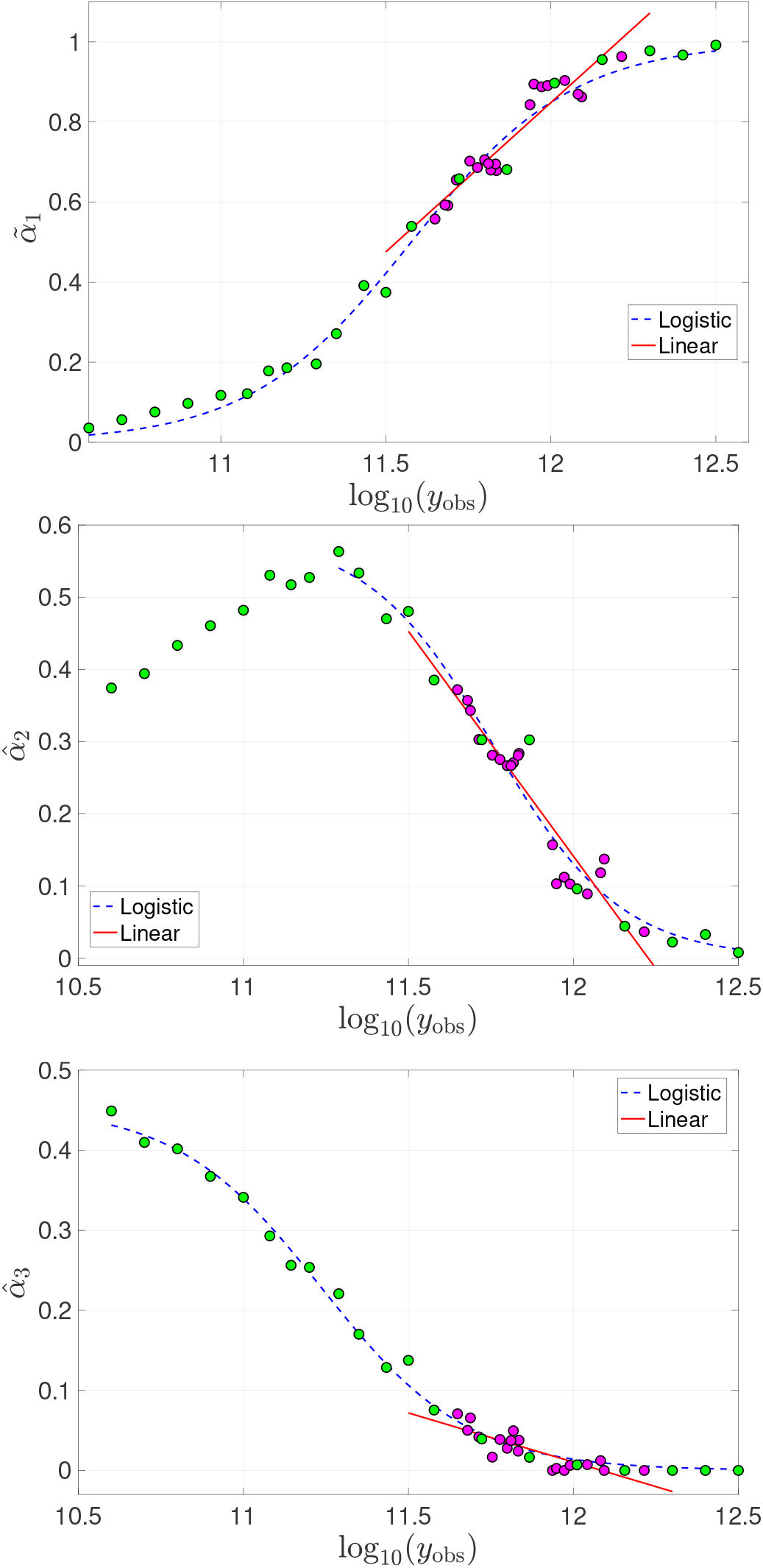
Relationship between the observed values and the estimated initial probabilities. Logistic and linear relationships between log_10_ (*y*_obs_) and 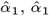, and 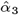.

To further investigate the relationship between log_10_(*y*_obs_) and 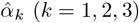, we added virtual observations (green disks) to extrapolate beyond the patient data over the range [10.6, 12.5]. We then fitted logistic curves (blue dashed) to the full set of points for 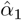 and 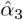. For 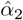, the data exhibit a less clear trend, so we restricted the logistic regression to the subset of points where the relationship starts to decrease.

The logistic regression functions for 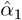 and 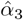 are respectively given by

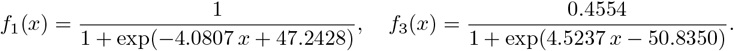

We note that if the range were extended below 10.6, it is highly likely that the relationship for 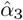 would take a similar shape to that of 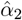, but this extension has been omitted here.

## Data Availability

All relevant data are within the manuscript and its Supporting Information files.

## Acknowledgments

Sylvie Vande Velde thanks the F.R.S.-FNRS for support through a Télévie grant. Sophie Hautphenne acknowledges support from the Australian Research Council (ARC) through Discovery Project DP200101281. Céline Engelbeen thanks the F.R.S.-FNRS and WBI for partial support of two research stays at The University of Melbourne to work on this project, and the ASEM-DUO programme for co-funding the second stay. Ilaria Pagani and David Ross thank the patients, investigators, study coordinators, and ALLG staff who provided the samples and clinical data for the study in [21], and acknowledge the ALLG as the sponsor of the TIDEL-II clinical trial. The authors thank Mingqi He for fruitful discussions on ABC and importance sampling, and Jacques Louis for his creative talent in designing Fig 1 and Fig 2.

